# Beyond target-decoy competition: stable validation of peptide and protein identifications in mass spectrometry-based discovery proteomics

**DOI:** 10.1101/765057

**Authors:** Yohann Couté, Christophe Bruley, Thomas Burger

**Affiliations:** Univ. Grenoble Alpes, CNRS, CEA, INSERM, IRIG, BGE, F-38000 Grenoble, France

## Abstract

In bottom-up discovery proteomics, target-decoy competition (TDC) is the most popular method for false discovery rate (FDR) control. Despite unquestionable statistical foundations, this method has drawbacks, including its hitherto unknown intrinsic lack of stability *vis-à-vis* practical conditions of application. Although some consequences of this instability have already been empirically described, they may have been misinterpreted. This article provides evidence that TDC has become less reliable as the accuracy of modern mass spectrometers improved. We therefore propose to replace TDC by a totally different method to control the FDR at spectrum, peptide and protein levels, while benefiting from the theoretical guarantees of the Benjamini-Hochberg framework. As this method is simpler to use, faster to compute and more stable than TDC, we argue that it is better adapted to the standardization and throughput constraints of current proteomic platforms.

---

Most projects involving mass spectrometry (MS)-based discovery proteomics use data-dependent acquisition workflows in which tandem mass (MS/MS) spectra are produced from isolated peptides. Then, peptide identification is performed by database search engines which match the experimental spectra acquired with theoretical spectra derived from a list of protein sequences ^1^. The more the experimental spectrum resembles the theoretical spectrum, the higher the matching score. This methodology has been widely adopted, but it was soon recognized that it could lead to false positive identifications ^2^. Indeed, among the tremendous number of spectra generated by a peptide mixture prepared from a complex biological sample, at least a few of them are expected to match an erroneous sequence, by chance. To avoid corrupting the biological conclusions of the analysis, researchers have come to rely on statistical procedures to limit the False Discovery Proportion (FDP) - *i*.*e*. the proportion of mismatches among all the peptide spectrum matches (PSMs) which look correct. As this quality control problem is ubiquitous in science, statisticians have extensively studied it. The main conclusions of these studies (See ^3^ for a proteomic-oriented summary) are as follows: *(i)* Due to the random nature of the mismatches, it is impossible to precisely compute the FDP; *(ii)* However, it can be estimated, as an FDR (False Discovery Rate); *(iii)* Depending on the experiment, the FDR will provide a more or less accurate estimate of the FDP; *(iv)* Therefore, practitioners should carefully select the FDR methodology, and interpret its result cautiously, making an educated guess (*e*.*g*., like a political poll before an election).

Target-decoy competition (TDC) has emerged as the most popular method to estimate the FDP in MS-based discovery proteomics ^4^ Its success is a marker both of its conceptual simplicity and of its broad scope of application. The principle of TDC is to create artificial mismatches by searching a specific (“decoy”) database of random sequences which differ from the sequences of interest (present in the “target” database) and to organize a competition between target and decoy assignments. Under the so-called Equal Chance Assumption (or ECA, stating that target mismatches and decoy matches are equally likely ^4^), it is possible, for any given cut-off score, to estimate the number of target mismatches that will be validated. Like any other estimator, TDC-FDR can lead to inconsistent estimates if the theoretical assumptions on which it is based do not hold in practice. Notably, the quality of TDC-FDR is strictly linked to the ECA validity, *i*.*e*. the decoy’s capacity to adequately fool the database search engine. If it fools it too much, the TDC-FDR will overestimate the FDP; whereas if it is too unrealistic to fool the search engine, the FDP will be underestimated^5^. For this reason, decoy database construction and conditions of application have been extensively studied. Results from these studies indicate that: *(i)* the search engine must be compliant with TDC ^6^; *(ii)* In theory, the larger the decoy database, the more precise the mismatch score distribution ^7,8^; *(iii)* The decoys must respect the cleavage sites ^9^ to avoid systematic target matching regardless of spectrum quality; *(iv)* The influence of randomness in the construction of the decoy database can be counter-balanced by boosting strategies, leading to less volatile FDRs ^10^; *(v)* Decoy counting also has an influence ^8^. In addition to these restrictions, numerous parameters have been reported and discussed to control their relative importance ^11^. This extensive body of literature has notably contributed to installing the competition step of TDC as essential, and today, target-decoy searches without competition ^12,13^ are scarcely ever reported. Despite the wide acceptance of TDC, a series of letters from Bret Cooper ^14,15^ initiated a controversy regarding the observed downfall of TDC validation levels with data produced by high-resolution mass spectrometers. He provided experimental arguments to reject the idea that such downfall was simply a positive consequence of instrument evolution, leading to an increase in the numbers of peptides identified. Notably, he pointed out that very low-quality spectra incompatible with confident peptide identifications could be validated despite application of a stringent FDR cut-off. Moreover, as this phenomenon was observed with multiple widely-used search engines (Mascot, X!tandem and MS-GF+), he concluded that there was an “*inherent bias*” of “*peptide presumption*” (i.e., only peptides already listed in the target database could be identified). As this stance contradicted both empirical and theoretical evidence, a few articles were published arguing against this view ^16,17^ while others confirmed^18,19^, maintaining the *statu quo*.

However, Cooper’s observations can be reconciled with statistical theory. In fact, the correctness of any statistical estimate is only asymptotic: if the quality of the empirical model depicting the mismatches is improved (for instance, by increasing the size of the decoy database^7,8^ or by averaging a growing number of TDC-FDRs resulting from randomly generated decoy databases, in a boosting-like strategy ^10^), we should end-up with a series of estimates that theoretically converges towards the FDP. Although essential, this asymptotic property is unfortunately not sufficient for practitioners, who work with a finite number of decoy databases of finite size (classically, a single decoy database of the same size as the target database). In this context, even if TDC is *asymptotically unbiased* (*i.e*., no systematic difference between the FDP and the FDR) it could sometimes lead to inaccurate FDRs (*i.e*., over-conservative or anti-conservative estimates) due to excessive variance (*i.e*., extensive stochastic fluctuations).

In this article, we shed new light on Cooper’s observations, which reconcile opposing opinions: While we believe target and decoy searches can be used to accurately compute FDRs, we uphold his concerns by showing that, with state-of-the-art high-resolution instruments, the risk that the TDC strongly underestimates the FDP increases. We then describe a series of mathematical transformations of classical identification scores, to which the well-known Benjamini-Hochberg (BH) procedure ^20^ and its numerous variants ^21^ can be applied at spectrum, peptide and protein levels. This leads to an original and powerful framework that demonstrably controls the FDR without decoy databases. Altogether, the results presented demonstrate that making TDC-FDR compliant with instrument improvements requires unexpected efforts (careful implementation, fine-tuning, manual checks and computational time), whereas identification results from MS-based proteomics can be simply and accurately validated by applying alternative strategies.

## Experimental Section

### Sample preparation and nanoLC-MS/MS analyses

For this work, we used the data obtained with a quality control standard composed of *E. coli* digest, analyzed according to a standard protocol.

Briefly, competent *E. coli* DH5*α* cells transformed with pUC19 plasmid were grown at 37°C in Lysogeny broth (LB) medium containing carbenicillin before harvesting during exponential phase (OD_600_ ∼ 0.6). After centrifugation at 3’000 × *g* during 10 min, the pellet was washed 3 times with cold Phosphate Buffered Saline (PBS) before lysis of cells using Bugbuster Protein Extraction Reagent (Novagen) containing cOmplete™, Ethylene-diaminetetraacetic acid (EDTA)-free Protease Inhibitor Cocktail (Roche) and benzonase (Merck Millipore). After centrifugation at 3’000 × *g* during 30 min and at 4°C, the supernatant was recovered and the protein amount was measured, before protein solubilisation in Laemmli buffer.

Proteins were stacked in a single band in the top of a SDS-PAGE gel (4-12% NuPAGE, Life Technologies) and stained with Coomassie blue R-250 before in-gel digestion using modified trypsin (Promega, sequencing grade) as described in ^22^.

Resulting peptides were analyzed by online nanoliquid chromatography coupled to tandem MS (UltiMate 3000 and LTQ-Orbitrap Velos Pro, or UltiMate 3000 RSLCnano and Q-Exactive Plus, Thermo Scientific). The equivalent of 100 ng of starting protein material was used for each injection. Peptides were sampled on 300 *μ*m × 5 mm PepMap C18 pre-columns (Thermo Scientific) and separated on 75 *μ*m × 250 mm C18 columns (Reprosil-Pur 120 C18-AQ, Dr. Maisch HPLC GmBH, 3 *μ*m and 1.9 *μ*m porous spherical silica for respectively UltiMate 3000 and UltiMate 3000 RSLCnano). The nanoLC method consisted of a linear 60-min gradient ranging from 5.1% to 41% of acetonitrile in 0.1% formic acid.

For LTQ-Orbitrap Velos Pro analyses, the spray voltage was set at 1.5 kV and the heated capillary was adjusted to 200°C. Survey full-scan MS spectra (*m/z* between 400 and 1600) were acquired with a resolution of 60’000 at *m/z* of 400 after the accumulation of 10^6^ ions (maximum filling time 500 ms). The twenty most intense ions from the preview survey scan delivered by the Orbitrap were fragmented by collision-induced dissociation (collision energy 35%) in the LTQ after accumulation of 10^4^ ions (maximum filling time 100 ms). MS and MS/MS data were acquired using the software Xcalibur (Thermo Scientific). For Q-Exactive Plus analyses, the spray voltage was set at 1.5 kV and the heated capillary was adjusted to 250°C. Survey full-scan MS spectra (*m/z* between 400 and 1600) were acquired with a resolution of 60’000 at *m/z* of 400 after the accumulation of 10^6^ ions (maximum filling time 200 ms). The ten most intense ions were fragmented by higher-energy collisional dissociation (normalized collision energy 30%) after accumulation of 10^5^ ions (maximum filling time 50 ms) and spectra were acquired with a resolution of 15’000 at *m/z* of 400. MS and MS/MS data were acquired using the software Xcalibur (Thermo Scientific).

### MS data analysis

Data were processed automatically using Mascot Distiller software (version 2.6, Matrix Science). Peptides and proteins were identified using Mascot (version 2.6) through concomitant searches against *Escherichia coli* K12 reference proteome (20180727 version downloaded from UniProt), and/or custom made decoy databases (reversed or shuffled sequences - see below). Trypsin/P was chosen as the enzyme and 2 missed cleavages were allowed. Precursor and fragment mass error tolerance has been variably adjusted as described in the manuscript. Peptide modifications allowed during the search were: carbamidomethylation (C, fixed), acetyl (Protein N-ter, variable) and oxidation (M, variable). Proline software ^23^ was used to filter the results: conservation of rank 1 peptide-spectrum match (PSM) and single PSM per query. 1% FDR control was performed with various methods, as described in the Results section. Precisions regarding the choice of the score (individualized vs. contextualized) as well as of alternative search engines can be found in Supporting Information S2.3 and S4.1.

### Decoy database generation

For classical TDC experiments Figures 1 and 2), we used the following procedure: The target database was reversed by using the Perl script `decoy.pl`) supplied with Mascot software and the generated decoy database was appended to the target one before concatenated search. From our observations, slightly different procedures shuffled vs. reversed, accounting for tryptic cleavage site, etc.) yields similar results, which concurs with prior knowledge ^11^.

**Figure 1:**
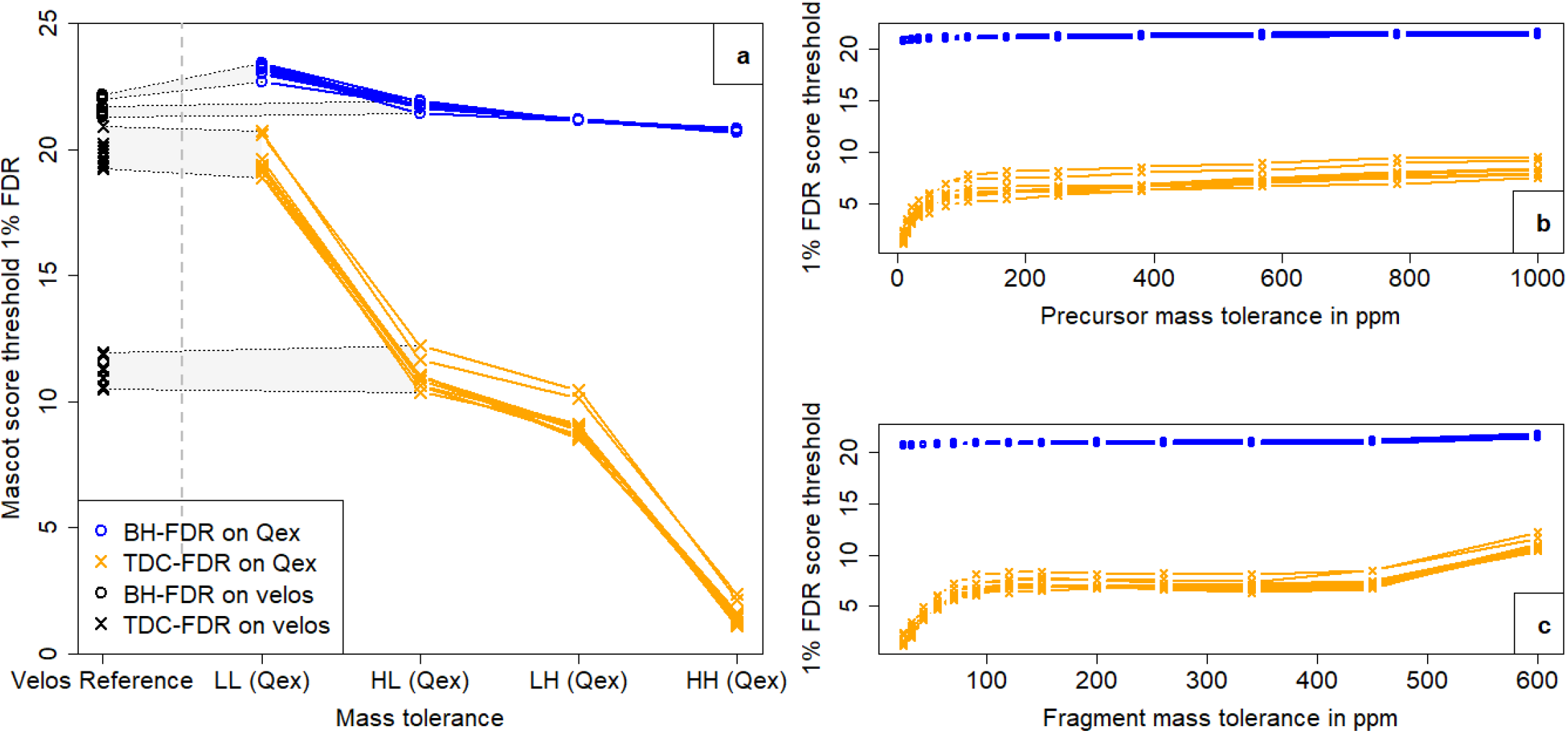
Score thresholds obtained when applying TDC (Orange) and BH (Blue) filtering at an FDR of 1%, as a function of the search engine mass tolerance parameters, for 10 samples analyzed with a Q-Exactive Plus (Qex) instrument. (**a**) Precursor and fragment mass tolerances were tuned to the LL, LH, HL and HH settings: LL assumes the MS and MS/MS data were acquired at low resolutions for the precursor and fragment masses (1 Da and 0.6 Da, respectively); HL uses mass tolerances of 10 ppm and 0.6 Da, respectively; LH uses mass tolerances of 1 Da and 25 mmu; and finally, HH uses mass tolerances of 10 ppm and 25 mmu (which corresponds to classical parameters for database searches performed with Qex data). The black lines encompass thresholds resulting from similar analyses performed on an LTQ-Orbitrap Velos Pro (Velos) with LL and HL settings. (**b**) Refined analysis of the FDR threshold s sensitivity to precursor mass tolerance tuning (Qex data, fragment tolerance = 25 mmu). (**c**) Refined analysis of the FDR threshold’s sensitivity to fragment mass tolerance tuning (Qex data, precursor tolerance = 10 ppm).

**Figure 2:**
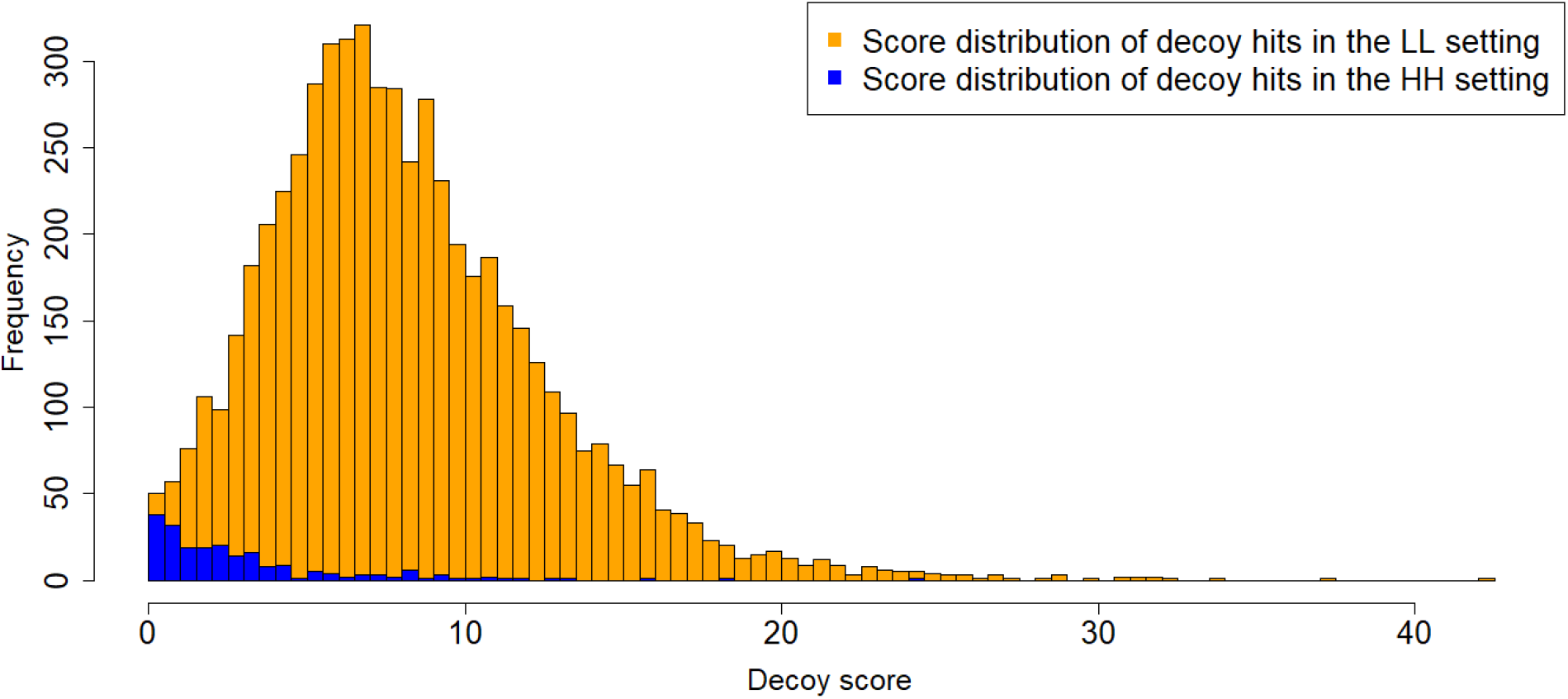
Distributions of scores for PSMs identified in the decoy database using the LL (orange) and HH (blue) settings. Data correspond to replicate 1, Qex analysis. After con-comitant searches in target and decoy databases using LL and HH parameters, the scores for the PSMs identified only in the decoy database, and without FDR filtering, were collected and represented as histograms (x-axis: score, bin 0.5; y-axis: frequency).

To compute the Empirical Null (EN) FDRs ^24^ shown in Figure 3, we relied on the model provided by a shuffled database not used in competition with the target database ^12,13^ combined with a boosting strategy ^10^ (*i.e*. a procedure averaging the FDR estimates from multiple shuffled decoy databases). For this study, we used 10 shuffled databases, each with a length equal to that of the target database, and produced by a shuffling procedure which respected cleavage sites to maintain the precursor mass distribution ^9^ R code available in Supporting Information S1.1).

**Figure 3:**
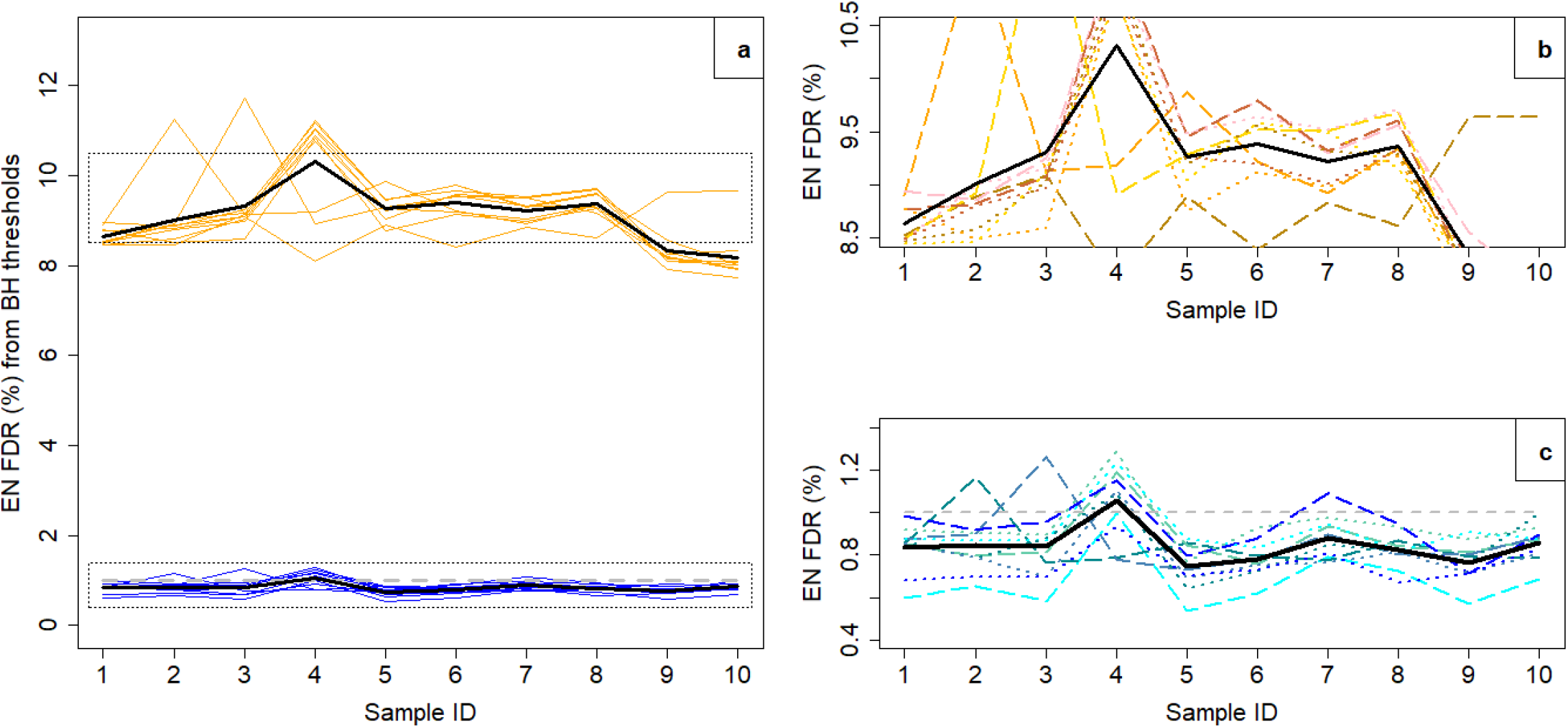
(**a**) Empirical Null (EN) FDRs computed for each replicate matched against 10 randomly generated (shuffled) decoy databases, according to its 1% FDR validation score threshold (gray dashed line), computed by applying BH (blue) and TDC (orange) methodologies; The black continuous lines show the average of the 10 EN FDRs (boosted estimate). (**b**) and (**c**) Zooms of the two framed areas in (**a**), with different shades of blue (resp. orange) and of line types (dot or dash) for better shuffle discrimination.

## Results and Discussion

### TDC and Benjamini-Hochberg procedures yield different FDRs

To counteract the the drop-off in validation levels observed with modern MS instruments ^14,15^, we suspected that the Benjamini-Hochberg (BH) approach to FDR control ^20^ could be an interesting alternative. Discrepancies between BH and TDC have already been reported ^25,26^, but unfortunately direct quantitative comparisons in terms of bias (*i.e*., systematic error) and variance (*i.e*., lack of stability) are impossible. ^27^ Such comparisons would require the FDP to be precisely known, although it is not accessible, even when using a controlled dataset. Thus, to better grasp the possible differences in behavior between FDRs estimated by the TDC and BH approaches, we reproduced Cooper’s experiment with the following extensions: First, both FDRs were computed simultaneously on the same datasets; Second, the settings for many experimental parameters were varied: we used multiple analytical replicates, different instruments, as well as various combinations of precursor and fragment mass tolerance sets.

To compute a BH-FDR, the search engine must provide PSM scores which can be related to p-values. Fortunately, numerous state-of-the-art search engines do so ^28^ : For instance, Mascot provides scores in the form

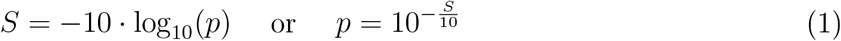

where *p* is a p-value. Andromeda provides a similar calculation, although the score is not directly accessible as it is only an intermediate computation (see Supporting Information S4.1). PepProbe, InsPecT and MyriMatch directly provide p-values as scores, and SEQUEST scores can be transformed into p-values through the application of dedicated wrappers *e.g*. ^25,26^. In theory, p-values always distribute uniformly across the [0,1] interval. However, when they do not in practice, they are termed *ill-calibrated or miscalibrated*. As miscalibration can affect BH estimates, potentially making them incorrect, tools have been developed to check or improve the p-value calibration (see Supporting Information S1.2).

We applied both TDC and BH methodologies to results acquired with a Q-Exactive Plus instrument on ten analytical replicates of an *E. coli* lysate. MS and MS/MS spectra were acquired at relatively high resolutions (70,000 and 17,500 at *m/z* of 200, respectively). We used Mascot to run a TDC search (see Experimental Section) and we considered four combinations of mass tolerance tuning at the precursor and fragment levels: LL, HL, LH and HH (where L stands for low precision or large tolerance, and H for high precision or narrow tolerance), the final combination (HH) corresponds to the tolerance levels generally used on our platform for Q-Exactive data analysis. In parallel, we ran target-only searches using the same parameters. Scores were then converted into p-values using Eq. 1, and the calibration of the resulting p-values was assessed as reported in Supporting Information S2.1. Finally, the classical BH step-up procedure for p-value adjustment was applied (see Data and code availability).

Figure 1A shows the score thresholds obtained at 1% FDR as a function of the mass tolerance combinations applied (see Supporting Information S2.2 for numerical values). At first glance, the drop-off in validation scores is obvious with TDC, whereas it is almost immaterial with BH. To avoid drawing sweeping conclusions on this impression, we performed complementary experiments. First, to better capture the influence of mass tolerance tuning, starting from the HH setting, we progressively extended the mass tolerance range, either for the precursor (Figure 1B) or for fragment masses (Figure 1C). The trends observed in these analyses support the one shown in Figure 1A. Moreover, they confirm that tolerances at precursor and fragment levels have a greater influence on threshold scores determined using TDC compared to BH.

Second, to confirm that the results obtained by reducing the mass tolerance in the search parameters mimics results obtained with lower-resolution instruments, we analyzed another batch of ten analytical replicates of the *E. coli* lysate submitted to MS/MS on a LTQ - Orbitrap Velos Pro with mixed resolutions: high resolution in MS mode (Orbitrap analysis) and lower resolution in MS/MS mode (linear ion trap analysis). Database searches were performed using the HL and LL tuning paramaters, and FDR thresholds were computed as above. Interestingly, the matching between the results from Q-Exactive and LTQ-Orbitrap data obtained using the same thresholds was excellent. This result justifies our methodology: from an FDR viewpoint, switching to analysis of a lower-resolution dataset using an appropriately-tuned search engine, or retaining the higher-resolution data while substantially increasing the mass tolerances, produces roughly equivalent outputs.

Therefore, we can interpret the different database search sets as surrogates for the recent improvements to instrumental capabilities: when TDC was first applied to data from low-resolution instruments, TDC and BH produced roughly similar results in terms of score cut-off to reach 1% FDR. Since then, the resolution of MS instruments has progressively increased, and now TDC and BH diverge considerably when applied to the same lists of putative PSMs, so that at least one of them (but possibly both) yields an incorrect FDR control.

### BH is more stable than TDC

Based on the results presented in Figure 1, BH appears more stable than TDC, both within and between experimental settings: First, for each tuning taken individually, the TDC threshold was less stable than its BH counterpart, as the set of ten cut-off scores was more dispersed with TDC. Second, depending on the mass tolerances applied when performing database searches, the TDC threshold on the Mascot score varied from 1.11 to 20.73 (with HH and LL settings, respectively), whereas its BH counterpart was more stable (between 20.67 and 23.43).

A part of the TDC’s instability (relative to BH) can be explained by the random nature of decoy sequence generation^10^, regardless of the search engine used. However, at first glance, there is no reason to assume that the remaining reported instability (notably the drop in score) is: (1) specifically linked to the TDC; (2) not a (much more important) bias issue. Consequently to point (1), it could make sense to question the algorithmic specificities of the search engine (here Mascot), as Cooper first did ^14^. Unfortunately, he then reported ^15^ similar pitfalls (at least for the precursor tolerance parameter) with X!tandem and MS-GF+ (formerly known as MS-GFDB). In addition, we discovered similar effects with Andromeda (Maxquant environment, see Supporting Information S2.3). Finally, our experiments demonstrate that not only the precursor mass tolerance set, but also, the fragment mass tolerance defined contribute to this effect. As for point (2), we hereafter demonstrate that even if the dependence on the search parameter is assumed to be a stability issue only (instead of a bias issue), it should nonetheless lead to legitimacy question TDC use.

### TDC instability can lead to anti-conservative FDRs

Naturally, more stable FDRs should be preferred, however less stable ones are not necessarily untrustworthy. As differently parametrized searches yield distinct putative PSM lists, different (correct) FDRs would make sense. However, the gap between BH and TDC cut-offs in the HH setting is alarming, as the PSMs identified in the target database before validation were the same with both methods. The very low cut-off scores obtained with TDC while using the HH setting led us to question the TDC procedure: Even if TDC is assumed to be unbiased, can its instability result in FDRs that are sometimes conservative (*e.g*. LL cases) and sometimes not (*e.g*. HH cases)? This context-dependent anti-conservativeness would clearly make use of the TDC approach less than reliable, as false discoveries would no longer be controlled.

To answer this question, we relied on the following rationale for FDR control: When a PSM list is filtered at 1% FDR, it does not mean that we accept 1% of additional poor matches in the result. On the contrary, it means that, even though all the validated PSMs apparently correspond to matches of sufficient quality, 1% of them are spurious (randomly distributed over the full range of scores, not necessarily matches with the lowest scores). Although it appears counter-intuitive, this property has already been empirically confirmed ^29^. Therefore, using only the HH setting and examining the list of PSMs with low scores validated by TDC is insightful to assess the conservativeness of this approach. We concretely did so with the results from the first analytical replicate, where 12209 PSMs were validated after applying TDC, leading to an expected number of mismatches at 1% FDR equal to 122. From the validated list, we therefore randomly selected 150 PSMs with a Mascot score < 10. The quality of the matches between theoretical and experimental spectra was obviously too low to yield confident identifications in the vast majority of cases (see Supporting Information S2.4). Based on this sample, the [1.61; 10[interval (which corresponds to only 635 PSMs, *i.e.* 5.53% of 12209) already contains more mismatches than the number expected for the entire dataset. This result is incompatible with 1% FDR validation, as random misidentification depicting apparently correct matches can also be expected at higher scores ^29^. In other words, on this dataset using the HH setting, the TDC FDR did not conservatively estimate the FDP.

These observations confirm those reported by Cooper. Although they are not sufficient to conclude on an intrinsic bias of TDC, they do confidently show that the drop-off in of validation cut-off scores is associated with, at least, an increased risk of anti-conservative FDR.

### Mechanistic explanation for TDC’s downfall

The above results contrast with the theoretical guarantees that have been published for TDC^7,30,31^. However, none of them account for the application of preliminary filters which reduce the number of decoy competitors, while Cooper’s controversy is rooted in such filters. This distinction may explain the discrepancy: Depending on the instrument’s accuracy when measuring the precursor mass, due to variable resolving powers, and assuming the search engine is tuned accordingly, a larger or smaller number of decoys are considered possible competitors for a given spectrum. Thus, MS data acquired with high-resolution mass spectrometers and analyzed using search engines in which narrow mass tolerances are *de facto* set leads to smaller numbers of decoy matches. Therefore, the ECA may no longer apply and the FDR may be underestimated. This situation can be experimentally observed while looking at the distributions of the PSMs identified in the decoy database using the LL and HH settings. Indeed, the number of decoy identifications, as well as their corresponding scores, were strongly decreased in the HH setting compared to the LL one (see Figure 2), and the very low number of decoy PSMs with the HH setting cannot provide an accurate FDR estimation.

As TDC appears to be inaccurate when the number of decoy challengers is too small, one straightforward solution would be to enlarge the database in proportion. Although the link between mass tolerance and decoy database size can be efficiently exploited to limit the computational cost of repeated searches^32^, it was, unfortunately, inefficient in our case (see Supporting Information S4.2).

### Decoy-based empirical null estimation and BH FDRs are consistent

An alternative means to investigate the above mechanistic intuition is to build a strategy in which the quality of decoy matches is preserved, regardless of the stringency of the database’s preliminary search filters; and then to examine how it estimates the FDR. The statistical theory of *empirical null estimation*^24^ is based on making false discoveries look like true ones. Even though this theory has already been applied in various proteomics contexts (giving rise to *entrapment methods*^27,33,37^), it provided us with a nice framework to elaborate on. Surprisingly, we found that a fair (but nevertheless unstable) empirical estimate of the null distribution could be obtained by performing a separate decoy search, without competition^12, 13^. To address the lack of stability, we averaged 10 repeated estimations, each based on a different shuffled database ^10^ (see Experimental Section). For this reason, for each of the 10 sample replicates of *E. coli* lysate analysed with the Q-Exactive Plus,Figure 3 shows the 10 empirical null (EN) FDRs corresponding to the cut-off scores obtained using BH and TDC 1% FDR filtering (HH setting).

The difference is striking: although BH thresholds (Mascot scores between 20.67 and 20.84) produced EN FDRs slightly below 1% (between 0.53% and 1.29%, with an average ≈0.84%), those obtained with TDC (scores between 1.11 and 2.37) led to 10-fold larger EN FDRs (between 7.72% and 11.71%, with an average ≈9.1%). This result is insightful for three reasons: *(i)* The fact that two orthogonal methods to compute an FDR (namely BH and EN) provided concurring results is an evidence supporting their correctness; *(ii)* it confirms that TDC can lead to considerable FDR under-estimations if used inappropriately; *(iii)* it shows that in contrast to Cooper’s concerns, the concept of “peptide presumption” is not inherently biased, since by applying an appropriate decoy search strategy, it is possible to cope for the cut-off downfall and to recover coherent FDRs.

### Practical comparison of EN and BH approaches

To summarize, the EN approach implemented here essentially amounts to averaging multiple target-decoy searches without competition, and can be viewed as an improvement of a 12-years-old method ^12,13^ well-fitted in the proteomics landscape, yet outmoded. In contrast, the BH approach is mainly theoretically motivated, and even though it is double the age, it is scarcely used in proteomics. As both methods provide concurring estimates, the one to promote mainly depends on their respective conditions of applicability and ease of use.

For BH: If we compare the EN FDRs derived from the BH-thresholds on Figure 3 with the expectation (i.e. 1%), BH appears to provide a slight overestimation. This result is probably due to the previously described over-conservative property of the BH estimator ^38^. Moreover, as previously mentioned, BH requires that the p-values first be checked for calibration. Fortunately, many methods can be used to limit BH over-conservativeness, and applying the best one can be done concomitantly with the calibration assessment, by means of a simple visual tool ^21^ (see CP4P description in Supporting Information S1.2). Finally, the BH FDR is extremely rapidly computed and does not require any decoy database.

For EN: Figure 3 shows a high dependency on the different decoy databases: From one randomly generated version to another, the FDR estimated varies significantly. At first glance, FDRs around 1% seem slightly more stable than those around 10%. However, after normalization relative to the mean FDR value, it is actually the opposite that occurs (mean coefficient of variation of 13.90% around 1% EN FDR, versus 5.82% around 10% EN FDR). This observation can easily be explained: With lower FDR thresholds, fewer decoys passed the threshold, and as a result, the statistics were computed on smaller sample sizes, inherently more sensitive to randomization. This explains why it is necessary to average multiple FDRs with different randomly generated decoy searches, despite the additional computational and practical complexity. However, depending on the complexity of the experiment, the precise number of searches required to stabilize the FDR cannot be estimated and will require manual trials. Moreover, if the experimental design requires iterative filtering of the database search (*e.g*. multiple-pass identifications ^39^), it is possible that the same anti-conservativeness issue will arise as with the competition step, so that additional caution should be applied. Therefore, compared to the EN strategy, the BH procedure is appealing for its simplicity and stability.

### FDR control at peptide level using the BH procedure

The difficulty of inferring peptide- and protein-level knowledge from spectrum-level information, while applying quality control criteria, has been widely addressed in the literature^40,41^. However, to our knowledge, all available inference systems require a preliminary decoy search to propose a peptide- or protein-level FDR. Today, combining multiple levels of FDR control has become accepted standard good practice. We therefore propose a generic procedure to extend the BH-FDR approach to peptide and protein levels. Moreover, the proposed method is independent of the chosen inference rules (see Supporting Information S3.1). Hereafter, we assume that the inference rules selected unambiguously define which PSMs should be used in peptide scoring, as well as which peptides contribute to protein group scoring ^36,37,42^, and we focus on the scoring methods applied.

The most classical peptide scoring methods assume that each peptide is identified by the spectrum with the highest PSM score amongst the *Q* matching spectra ^35-37,43^. In this setting, it makes sense to define the peptide score as equal to the best PSM score ^35^. Formally, if the **PSM score** between peptide sequence **seq**_*i*_ and spectrum *q* is referred to as *S*_*iq*_, then, the **best-PSM score** can be defined as max_*q*∈⟦ 1,*Q*⟧_ *S*_*iq*_ where ⟦*·, ·* ⟧. denotes an integer interval. This score can potentially be used to compute a TDC-FDR, but not a BH-FDR. Indeed, its probabilistic counterpart cannot be well-calibrated (the minimum of several calibrated p-values is non-uniformly distributed, see Figure S1). Fortunately, according to the following proposition, it is possible to modify the best-PSM score by applying a formula akin to Šidák correction^44^ and thus to recover correct calibration:

#### Proposition 1

*Let S*_*1*_; …: ; *S*_*n*_ *be a set of n scores of the form S*_*ℓ*_*=* −10 log_10_*(p*_*ℓ*_*) (ℓ ∈* ⟦1, *n* ⟧) *where p*_ℓ_ *is realizations of n i.i.d*. ℝ_*+*_ *random variables, X*_1_,…, *X*_*n*_ *If X*_*ℓ*_ *∼* 𝒰[0, 1] ∀ *ℓ,then*,

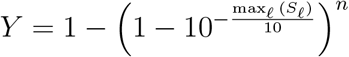

*uniformly distributes over the range* [0; 1].

*Proof:* See Supporting Information S3.2.

Therefore, (See Supporting Information S3.2 and S3.3 for the full derivations), the **peptide p-value** 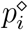 and **peptide score** 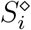 of peptide sequence **seq**_*i*_ can be defined as:

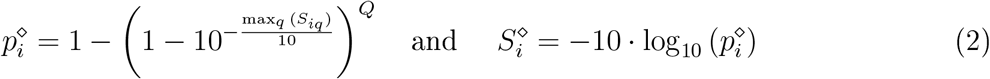

### FDR control at protein level using the BH procedure

To define protein-level scores and p-values, fragment matches for PSM scores were considered equivalent to what peptide matches are for protein scores. This equivalence led us to rely on Fisher’s test to define protein scores/p-values from the scores of the best subset of peptides. Similar approaches have frequently been investigated in the literature ^32,36,37,42^ and the full derivation is presented in Supporting Information S3.4. To the best of our knowledge, we are the first to discuss the adaptation of Fisher’s methodology from its original context (meta-analysis) to proteomics by explicitly considering (*i*) risks of anti-conservativeness due to dependent peptides (see Supporting Information S3.5); (*ii*) the impact of the poorly conclusive peptide-level evidence in an open-world assumption context (see Supporting Information S3.6). Finally, for a protein sequence **seq**_*π*_ identified by *K* specific peptides with scores 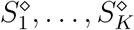, the **protein p-value** is defined as:

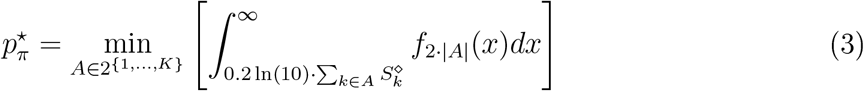

and the **protein score** as 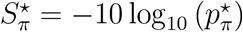, where: 2^{1,…,*K*}^ is the powerset of the set of *K* peptides identified; *A* is a peptide set with cardinality |*A*| ≤ *K*; and *f*_2·|*A*|_ is the density function of the *χ*^2^ distribution with 2 · |*A*| degrees of freedom. Although possibly obscure at first glance, 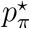 simply interprets as the p-value resulting from Fisher combined probability test applied to the subset of peptides which best explains the protein (see Supporting Information S3.4).

### Experimental assessment of BH FDR at peptide and protein levels

The ten replicate analyses of *E. coli* lysate were validated at 1% FDR by applying the BH procedure to the PSM, peptide and protein scores. To do so, only a target database search was necessary. However, and because it delivered a striking illustration of the capacity of the proposed framework to distinguish false identifications, we introduced shuffled sequences in the searched database to assess the results (see Experimental Section). We considered a challenging scenario where the number of decoys was set to five times the number of target sequences. Table 1 summarizes the average (across the 10 replicates) cut-off scores as well as the average counts for validated PSMs, peptides and proteins in both target and fivefold shuffled databases (see Table S2). Although the corresponding proportions must not be interpreted as FDRs, it is interesting to discuss them: First, despite the fivefold decoy over-representation, each of the three validation levels (PSM, peptide or protein) taken individually was sufficient to provide a decoy ratio below the FDP expectation of 1% at any level. Second, the three validation strategies provided broadly concurring filters and validated protein list sizes. Third, some discrepancies between the three validation strategies exist (for instance, when filtering at protein level, PSMs with low score are validated because they belong to proteins which are confirmed by other high scoring peptides), leaving room to refine validation with appropriate multi-level filters, as discussed below.

**Table 1:** Average (across the 10 *E. coli* replicates) minimum score and PSM, peptide and protein counts assigned as target and decoy in the raw dataset (No validation), as well as after validation by one of the three following rules: 1% BH-FDR at PSM level (PSM validation), 1% BH-FDR at peptide level (Peptide validation) and 1% BH-FDR at protein level (Protein validation).

|  | No validation |  |  | PSM validation |  |  | Peptide validation |  |  | Protein validation |  |  |
| --- | --- | --- | --- | --- | --- | --- | --- | --- | --- | --- | --- | --- |
|  | PSM | Pep. | Prot. | PSMs | Pep. | Prot. | PSM | Pep. | Prot. | PSM | Pep. | Prot. |
| Min score | 0.002 | 0.002 | 0.002 | 21 | 19.676 | 21.187 | 0.023 | 20.963 | 21.163 | 0.013 | 0.021 | 22.534 |
| #Targets | 12020.2 | 9351.3 | 1466.5 | 10233.6 | 8180.3 | 1297.7 | 10736.2 | 8159.5 | 1297.9 | 11836.6 | 9169.4 | 1294.6 |
| #Decoys | 873.3 | 817.2 | 786 | 11.5 | 10.9 | 10.7 | 11.1 | 10.2 | 10 | 13.8 | 12 | 9.2 |

## Conclusions

This work sheds new light on a crucial step in bottom-up proteomics experiments: the validation of identification results. First, it illustrates that the TDC and BH estimates of the FDP have progressively diverged as MS accuracy has improved. Our results demonstrated that this divergence originated in the TDC’s lack of stability with respect to the precursor and fragment mass tolerances set during database searches. Although this lack of stability can be partially counteracted by suppressing the competition step of TDC^12,13^, the instability induced by the random generation of decoy sequences remains^10^. Therefore, even though target-decoy strategies can be refined to partially cope with this instability, the results are not satisfactory as these strategies: *(i)* are not as stable as BH, in particular at lower FDRs; *(ii)* are more complex to organize (implementation, computational cost) and require additional manual checks; *(iii)* do not provide any guarantee of reliable FDR estimates in the future (on datasets acquired with even higher resolution next generation instruments for which narrower mass tolerances can be expected; or with pipeline modifications that change the number of target and decoy candidates).

Second, this work provides new peptide and protein scores which demonstrably respect the calibration conditions of the BH procedure. Indeed, implementing BH-FDR at PSM-, peptide- and protein-level is straightforward (see Data & code availability) and its practical use within a preexisting platform pipeline requires no precise tuning. Moreover, our results highlighted that, despite slightly different behavior, any of these scores alone is sufficient to conservatively validate a proteomics dataset at PSM, peptide and protein levels. This finding suggests that various strategies could be developed to comply with different objectives: If the expected output is a protein list, then it is probably most appropriate to control the FDR at protein-level. However, in studies seeking to refine discrimination between proteoforms sharing many subsequences, it may be more relevant to validate at peptide level. Finally, when quantifying proteins, extracting the ion current for misidentified spectra produces erroneous results, making validation at PSM level necessary. Beyond these considerations, acting at different levels of filtering may also improve the quality of the validated identifications, although this assertion requires further investigation. For example, multiple FDRs are classically used sequentially, following the inference process (starting at PSM level and ending at protein level); using a reverse order or parallel filtering may also be of interest to preserve the calibration necessary to the BH procedure.

Based on these results, we propose an overhaul of how FDR is estimated in discovery proteomics using database searching and suggest replacing TDC by BH-FDR. Nevertheless, as a theoretical research field, TDC remains of interest. The original idea proposed by Elias and Gygi ^4^ has stimulated the field of theoretical biostatistics and led to the idea that simulating null tests from the data (termed *knockoffs* instead of decoys) could produce efficient alternatives to BH procedures, which demonstrably control the FDR^4^. Transferring these theoretical results into biostatistics routines that can be applied on a daily basis still requires some investigation ^46, 47^. However, they will hopefully contribute to computational proteomics in the future, as an example of an interdisciplinary virtuous circle.

## Data and code availability

Implementing BH-FDR at PSM-, peptide- and protein-level is straightforward. First, if the scores of all the PSMs indicating a given peptide sequence are stored as a vector, `psm.scores`, then, the peptide p-value `pep.pval` and peptide score `pep.score` can be determined by applying the following `R` code:

~~~
library(Rmpfr) # to avoid roundings in p-values
psm.pvals <-mpfr(10**(-psm.scores/10), 128)
pep.pval <-1-(1-min(psm.pvals))^ length(psm.pvals)
pep.score <--10*log10(pep.pval)
~~~

Then, the protein score `prot.score` and p-value `prot.pval` for a protein for which peptide scores are stored in a vector `pep.scores` can be computed using the following code:

~~~
pep.scores=sort(pep.scores, decreasing=T)
nb.pep=length(pep.scores)
pep.cumscore=cumsum(pep.scores)*log(10)/5
tmp.scores=rep(0,nb.pep)
for(j in 1:nb.pep){
tmp.scores[j]=pchisq(pep.cumscore[j],2*j,lower.tail=F,log.p=T)
tmp.scores[j]=tmp.scores[j]/(−0.1*log(10))
}
prot.score= max(tmp.scores)
prot.pval= 10**(-prot.score/10)
~~~

Once the peptide scores and protein scores are available alongside the PSM scores provided by the search engine, the BH procedure can simply be run by applying the `p.adjust() R` function (base function). All these scores (and the BH procedure) are also implemented in Proline software ^23^, written in Java/Scala, so that any proteomics data analyst can use them whatever their coding skills. The mass spectrometry proteomics data have been deposited to the ProteomeXchange Consortium via the PRIDE ^48^ partner repository with the dataset identifier PXD016669 and 10.6019/PXD016669 (https://www.ebi.ac.uk/pride/, Username, Password: eA7lMPa7). A portable version of Proline software can be downloaded at ftp://ftp.cea.fr/pub/edyp/Proline/BH-FDR/. For each of the ten *E. Coli* replicates, it contains the search results from Mascot after the database search in the target and the fivefold decoy databases.

## Acknowledgments

This work was supported by grants from the French National Research Agency: ProFI project (ANR-10-INBS-08), GRAL project (ANR-10-LABX-49-01), Data@UGA and SYMER projects (ANR-15-IDEX-02) as well as MIAI @ Grenoble 718 Alpes (ANR-19-P3IA-0003). The authors are grateful to EDyP platform engineers, who performed the various quality control analyses used in this work, and notably to Alexandra Kraut for the sample preparation, as well as the ProFI developers of Proline software (http://www.profiproteornics.fr/proline/), which was used to perform all the data analyses presented. Finally, the authors thanks the anonymous reviewers for their rich and constructive comments.

## Author contributions and competing interests

Y.C. proposed the experimental methods and processed the data. C.B. developed the software environment to perform the experiments and implemented the proposed methodology. T.B. proposed the theoretical framework and drafted the manuscript. All authors contributed to the manuscript and approved its final version. The authors declare no competing interests.

## Supporting Information Available

- PDF document containing Supporting tools (S1), Supporting results (S2), Supporting methods (S3) and Supporting discussion (S4)
- Zip folder “spectra.zip” containing 150 spectra in PNG format (S5)

