## Supplemental Material for "Beyond target-decoy competition: stable validation of peptide and protein identifications in mass spectrometry-based discovery proteomics"

September 8, 2020

### S1 Supporting tools

#### S1.1 R code to shuffle the decoy database

```
library(stringr)
library(stringi)
library(Peptides)
options(warn=2)

##############################
### User defined parameters
##############################

# working directory:  must be completed and uncommented
# WARNING!!! - \ should be replaced by / when copy-pasting from windows
# setwd("C: XXXX")

# the input fasta file MUST only contain the target proteins
file <- scan("UP_K12_RefProt_D_20180727_target_only.fasta",
              what="character", sep="\n")

# Define which peptide to shuffle, or not
minimum.shuffled.Mass <- 400
maximum.shuffled.Mass <- 4800
minimum.length.ifnomass <- 5 #impossible to compute molecular weight if amino acid X occurs
assign("list.Of.Prec.Ions", NULL, envir = .GlobalEnv)

# Number of randomization
# If tuned to N, then, the decoy database is N times larger than the target one
# WARNING!!! doing so will break the Equal Chance Assumption
number.of.shuffling <- 1

# the name of the ouput file -- modify the first argument according to the database name
# WARNING!!! If the file already exist, delete it, or the output will be concatenated!
output.file <- paste("UP_K12_RefProt_D_20180727_target",
                     "_rand", number.of.shuffling, ".fasta", sep="")

###############################
### atomic functions
###############################

# finds tryspic cleavage site
getPeptides <- function(the.protein){
  cleavage <- str_locate_all(the.protein, "K|R")[[1]][,1]
  nbCleavage <- length(cleavage)
  if((nbCleavage ==0) || ( (nbCleavage ==1) &&
                         (cleavage==str_length(the.protein))
                         ) ){
    pep.bounds <- matrix(c(1, str_length(the.protein)), ncol=2)
  } else{
    npep <- nbCleavage+1
    pep.bounds <- matrix(rep(0,2*npep), ncol=2)
    pep.bounds[1,1] <- 1
    pep.bounds[2:npep,1] <-cleavage+1
    pep.bounds[1:(npep-1),2] <-cleavage
    pep.bounds[npep,2] <- str_length(the.protein)
    if(pep.bounds[npep-1,2] == str_length(the.protein)){ #no last peptide
      pep.bounds <- pep.bounds[-npep,]
    }
```

```
  }
  return(pep.bounds)
}

# shuffle a peptide if in the mass window
pepShuff <- function(the.peptide, minM=800, maxM=48000, minL=7){
  if(
    (dim(str_locate_all(the.peptide, "X")[[1]])[1] != 0) ||
    (dim(str_locate_all(the.peptide, "U")[[1]])[1] != 0)
    ){
    mass <- 0
  } else{
    mass <- mw(the.peptide) # compute molecular weight from Peptide package
    assign("list.Of.Prec.Ions", c(list.Of.Prec.Ions, mass), envir = .GlobalEnv)
  }
  peplen <- str_length(the.peptide)
  if( ( (mass>=minM) && (mass<=maxM) ) || ( (mass==0) && (peplen>=minL) ) ) {
    Cter <- str_sub(the.peptide, start=peplen, end=peplen)
    if( (Cter == "K") || ( Cter == "R") ){
      b <- peplen-1
      shuf <- paste(stri_rand_shuffle(str_sub(the.peptide, 1,b)), Cter, sep="")
    } else{ # last peptide of the proteins, does not end by a cleavage site
      shuf <- stri_rand_shuffle(the.peptide)
    }

  } else {
    shuf <- the.peptide
  }
  return(shuf)
}

proteinShuffler <- function(the.protein){
  peplist <- getPeptides(the.protein)
  nbpep <- dim(peplist)[1]
  tmp.seq <- ""
  for(i in 1:nbpep){
    the.peptide<- str_sub(the.protein,
                          start=(getPeptides(the.protein)[i,1]),
                          end=(getPeptides(the.protein)[i,2]))
    newpep <- pepShuff(the.peptide,
                       minM=minimum.shuffled.Mass,
                       maxM=maximum.shuffled.Mass,
                       minL=minimum.length.ifnomass)
    tmp.seq <- paste(tmp.seq, newpep, sep="")
  }
  return(tmp.seq)
}

# create shuffled database
createShuffledSeq <- function(list.tar){
  list.shuf <- list()
  for(i in 1:length(list.tar)){
    the.protein <- list.tar[[i]]
    randprot <- proteinShuffler(the.protein)
    list.shuf <- c(list.shuf, randprot)
  }
  return(list.shuf)
}
```

```
modifyListName <- function(list.name, j){
  list.name.decoy <- list()
  for(i in 1:length(list.name)){
    protname <- list.name[[i]]
    sp <- strsplit(protname,split='|', fixed=TRUE)
    l <- length(sp[[1]])
    nm <- paste(sp[[1]][1], "|",
            sp[[1]][2], "|", "###REV###RANDOM",j,"###",
                sp[[1]][3:l], sep="")
    list.name.decoy <- c(list.name.decoy, nm)
  }
  return(list.name.decoy)
}

### test atomic functions
# the.protein <- list.seq[[1]];
# the.peptide
# pepShuff(the.peptide)
# str_sub(the.peptide, 35,str_length(the.peptide))
# pepShuff(str_sub(the.peptide, 35,str_length(the.peptide)))
# the.protein
# proteinShuffler(the.protein)
# list.shuf <- createShuffledSeq(list.seq)
###############################################################

# count the number of target proteins and initialize the variables
prot.name <- rep(0,length(file))
for(i in 1:length(file)){
  if(substr(file[i], 1,4) == ">sp|"){
    prot.name[i] <- 1
  }
}
nb.prot <- sum(prot.name)
list.name <- list()
list.seq <- list()
tmp.seq <- "init"

# split the protein names and sequences, unsplit the sequences
for(i in 1:length(file)){
  if(prot.name[i] == 1){
    list.name <- c(list.name, file[i])
    if(i !=1){list.seq <- c(list.seq, tmp.seq)}
    tmp.seq <- ""
    } else{
      tmp.seq <- paste(tmp.seq, file[i], sep="")
    }
}
list.seq <- c(list.seq, tmp.seq)

# check the number of proteins and the corresponding list length
length(list.name)
length(list.seq)

for(i in 1:nb.prot){
  write(c(list.name[[i]],list.seq[[i]]), file=output.file, append=TRUE)
```

```
}
print("Pre-processing done and targets dealt with")
for(j in 1:number.of.shuffling){
  list.name.decoy <-modifyListName(list.name, j)
  #write(paste("XXX DECOYS",j,"XXX",sep=""), file=output.file, append=TRUE)
  list.decoys <- createShuffledSeq(list.seq)
  for(i in 1:nb.prot){
    write(c(list.name.decoy[[i]],list.decoys[[i]]), file=output.file, append=TRUE)
  }
  print(paste(j, "th randomization done and saved"))
}

hist(list.Of.Prec.Ions, breaks=100, xlim=c(0,8000),
     xlab="Theoretical precusor mass", main=" ")
```

#### S1.2 Calibration tools

Here, we survey the tools available to ensure BH procedure can confidently be applied to search engine outputs. This section is organized from the worst case (no p-value available), to the best one (well-calibrated p-values), and it ends with the presentation of CP4P (Calibration plot for proteomics [1]) a versatile tool that can be used to assess the quality of the p-value calibration, to partially improve it in case of ill-calibration, and to limit the well-known over-conservative behavior of BH procedure.

##### S1.2.1 Transforming scores into p-values

Few search engines do not provide any p-value while the provided score cannot be related to a p-value (e.g. Morpheus [2] or X!tandem [3]). However, as long as the scoring system used does not involve the PSM distribution, but on the contrary, solely relies on each spectrum sequence pair (see Supporting Information S4.1) it is possible to follow the recommendations of [4] to obtained calibrated p-values by means of a decoy database search. However, in absence of precisely described procedure in the article, many implementation details are left to the reader. Based on the recommendations of [5] (which notably highlights the importance of making a distinction between best-scoring and lower ranked decoy PSMs) we ended up with two different calibration procedures, both of which being apparently partly compliant[1] with [4]:

**Procedure 1**

1. Generate a decoy database of length $N$, $N$ being arbitrary large (the larger $N$, the better), and run the search engine on this decoy database (no target database involved)
2. Order all the queried PSMs with decreasing confidence: $\{S_1, \ldots, S_i, \ldots, S_m\}$ (where $m$ is the number of scored PSMs, $m \leq N \times \ell$ where $\ell$ is the number of spectra) and create a score to p-value conversion table by applying the following rule:

$$p\text{-value}(S_i) := i/m$$

3. For any new target database search (without decoy search), each PSM receives a p-value corresponding to that of the score $S_i$ it is the closest to in the conversion table (when the PSM score falls somewhere between $S_i$ and $S_{i+1}$, it is safer to consider the latter one, leading to a larger p-value, to enforce conservativeness).

Such approach however leads to two issues: First, some search engines do not grant access to all the PSMs, but only to the best scoring ones, so that it is impossible to build the right-hand side of the distribution; Second, the resulting p-values will need Šidák correction (see next subsection) before applying BH procedure. Alternatively, it is possible to define a procedure that is immune to both aforementioned drawbacks. Yet, as a counterpart, it will be only available for a given target database size:

[1] In [4], it is reported that "In practice, the decoy database is usually the same size as the target database; however, this is not necessary. Using a larger decoy database leads to more accurate p-value estimates at the expense of more computation.". However, from our experiments, Procedure 1 works to compute p-values, but cannot be straightforwardly extended to compute an FDR, unless the decoy database if of the same size as the target one; Conversely, when we applied Procedure 2 with a decoy database of different size, we obtained uncalibrated results.

**Procedure 2**

1. Generate a decoy database of length $N$ ($N$ being equal to the target database used), and run the search engine on this decoy database (no target database involved)

2. Order all rank-1 PSMs with decreasing confidence: $\{S_1, \ldots, S_i, \ldots, S_\ell\}$ (where $\ell$ is the number of spectra; the larger $\ell$, the more accurate the p-values) and create a score to p-value conversion table by applying the following rule:
$$p\text{-value}(S_i) := i/\ell$$

3. For any new target database of size $N$, perform a search without considering the decoy database, and assign to each PSM a p-value corresponding to that of the score $S_i$ it is the closest to in the conversion table (when the PSM score falls somewhere between $S_i$ and $S_{i+1}$, it is safer to consider the latter one, leading to a larger p-value, to enforce conservativeness).

Beside these two procedures, p-value calibration has recently gain interest, and a growing number of articles propose methods to convert search engine scores into well-calibrated p-values, either dedicated to a single search engine (e.g. [6, 7, 8] for Tides and SEQUEST algorithm related software), or applicable to many of them (e.g. [9]).

##### S1.2.2 Šidák correction may be needed in some cases

As outlined in [7], sometimes, an additional multiple testing correction is needed. The reason is the following: A statistical test is termed "calibrated" when the observations are tested against a distribution corresponding to that of the null hypothesis. Thus, the level of calibration correctness tightly relates to how the null hypothesis is defined. However, in our case, two null hypotheses are equally meaningful: First, from the search engine viewpoint, the null distribution is made of all the possible mismatches. However, the user seldom has access to this entire set of mismatch: Most of the time, only a subset containing the best target or decoy matches is provided; or ultimately, only the best-scoring PSM. Thus, from the user viewpoint, the null distribution is the distribution of the best-scoring mismatches (out of several possibilities). The thing is that the minimum of several uniform p-values (corresponding to the best-scoring peptide-spectrum mismatch) is not uniform, as illustrated on Figure S1. To cope for this [7], it is necessary to adjust the p-value of each PSM $i$ by accounting for the number $N_i$ of sequences that were tested, which is the purpose of Šidák correction:

$$p_i := 1 - (1 - p_i)^{N_i}$$

Importantly, search engines can either directly embed this Šidák correction, or not. Both definitions are acceptable, and both can be considered as leading to well-calibrated p-values. However, to apply BH procedure, it is necessary to have a uniform distribution for the scores of the best mismatches, so that if not embedded, this correction must be applied first.

Finally, let us mention the following warning: if the p-values are equal with or without Šidák correction, then it means $N_i = 1$. In other words, the preliminary filters on the database searched are so stringent that in practice, only a single sequence is eligible; which dramatically questions the identification pipeline, as broadly, one forces the identifications towards sequences of interest.

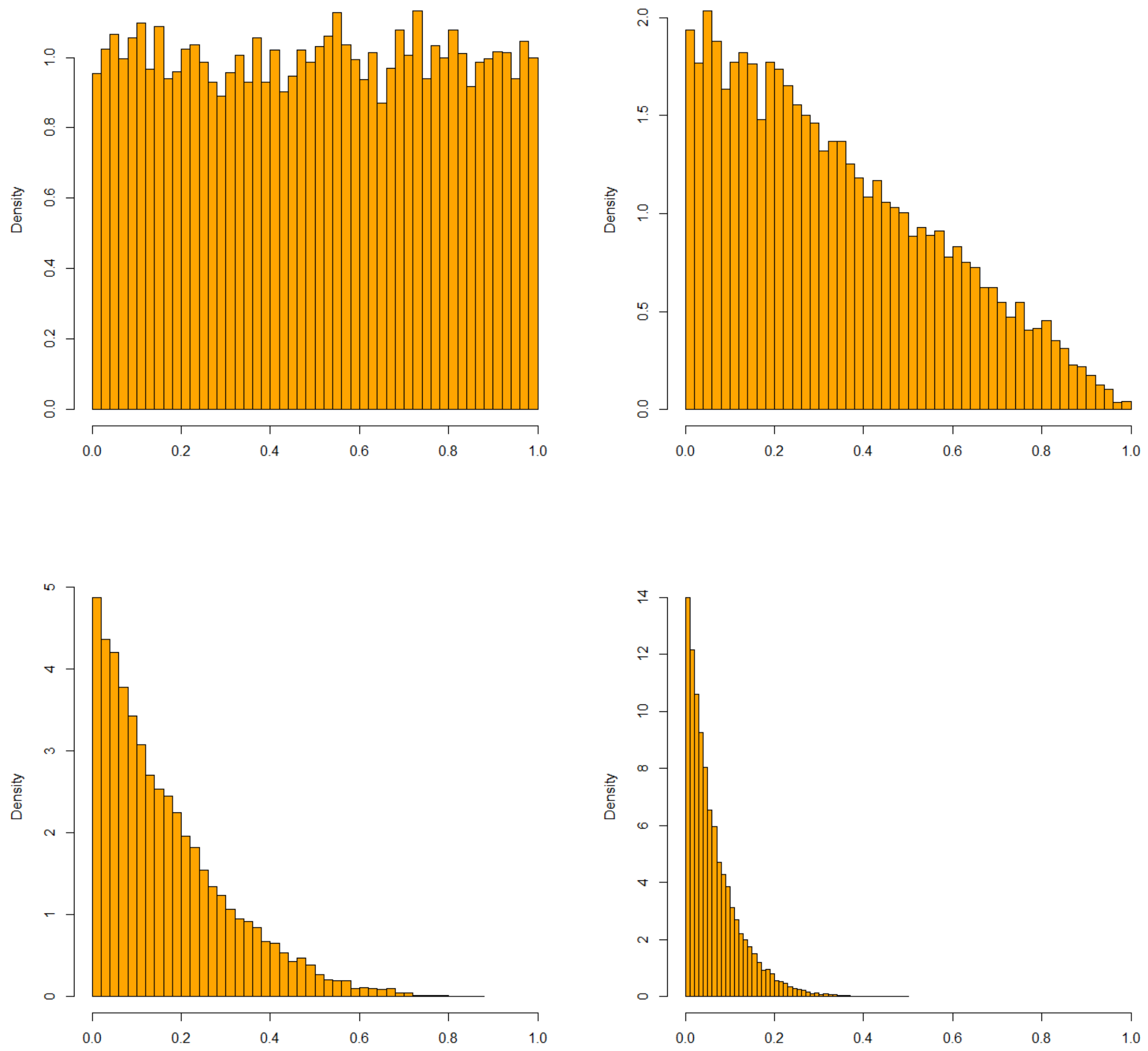

Figure S1: Histogram of 15,000 simulated p-values, after taking the minimum out of $N$ uniformly distributed samples, with $N = 1$ (upper left), $N = 2$ (upper right), $N = 5$ (lower left) and $N = 15$ (lower right): by taking the minimum p-values, one promotes small p-values with respect to large p-values, so that from a distribution with used to be uniform, one ends up with another distribution with is shifted to the left.

##### S1.2.3 Assessing the correct calibration of the p-values

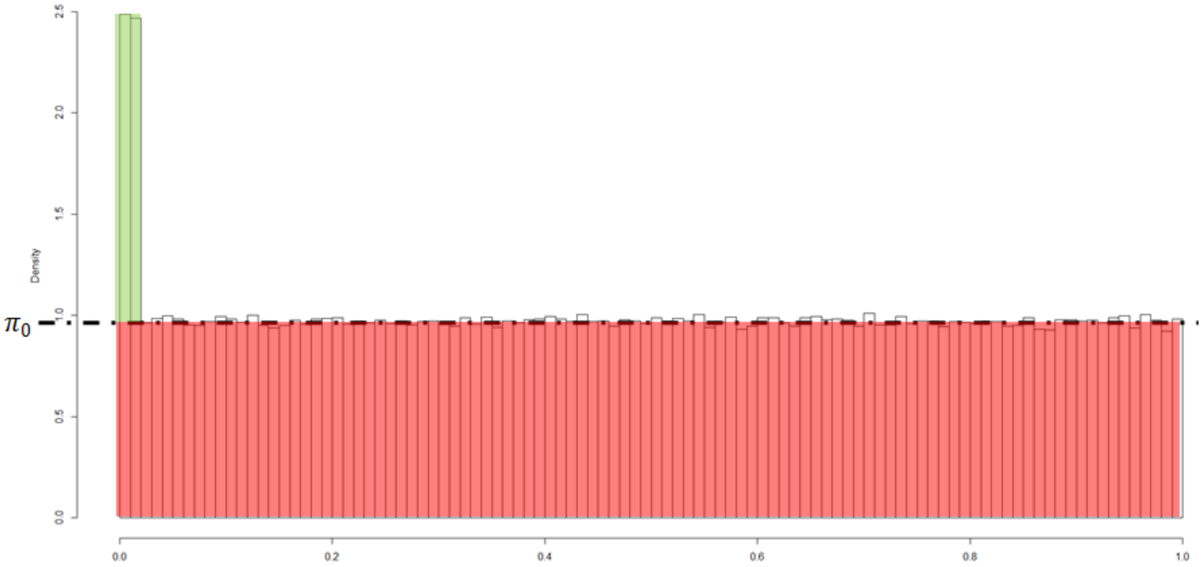

Figure S2: p-value histogram for a dataset where a proportion of $\pi_0$ false discoveries and of $1 - \pi_0$ true discoveries coexists (here $\pi_0 = 0.97$) – borrowed from [10].

Several methods are available to determine the extent to which p-values are calibrated. We can broadly divide them in two categories: those which have thrived on the proteomics context of extensive database search use; and those which root on statistical metrics. The former are more intuitive to proteomics practitioners but are often more hand-crafted too.

A first method [6] is to apply one of the two decoy-based procedures described above (depending on one assumes Šidák correction to be embedded or not), and to plot (in log-scale for better visibility) these reference p-values against the p-values to be checked (on the same decoy dataset). In case of correct calibration, the scatter plot should form a diagonal line with a unitary slope.

A second family of methods (also relying on database search engines) is referred to as *entrapment methods* [11, 12, 13, 14, 15, 16]. Conceptually, entrapment databases are essentially decoy databases, which are constructed in a specific way, and then searched in a specific way, so that the distinction between "decoy" and "entrapment" sequences is sometimes forgotten [12]. Moreover, their use has progressively shifted across the years: while they initially focused on p-value calibration [11], they have been extended to approximate the FDP so as to estimate the (FDR, FDP) difference [14], to finally serve as a new FDR computation method [15].

Based on more statistical assumptions, the easiest way to check the calibration of the p-values is to display their histogram [17] and to check visually the distribution shape: one should observe a rectangular histogram of hight $\pi_0$ (where $\pi_0 \leq 1$ depicts the proportion of null hypotheses), except for a peak on the left-hand side which depicts the discoveries with p-values outlying from the uniform distribution, see Figure S2. However, histograms tend to be unstable due to the binning, unless made of several thousands of p-values. To cope for this, it is possible to replace the p-value histogram by a cumulative distribution representation, as proposed in the early 80's by Scheweder and Spjøtvoll [18]. Let us by the way note that according to Yoav Benjamini [19], Scheweder and Spjøtvoll paper was at the root of BH procedures, published 13 years afterwards [20]. However, interpreting Scheweder and Spjøtvoll plot is not straightforward. This is why, we have recently adapted it to the proteomics context (and more precisely to the differential analysis problem), giving birth to the CP4P tool [1] that is presented below.

##### S1.2.4 How to deal with miscalibrated or ill-calibrated p-values

So far, we have only discussed how to assess the p-value calibration. If the p-values are perfectly calibrated, it is possible to directly proceed with BH procedure. However, in practice, it is possible to observed other situations. Broadly, we can consider three different scenarios:

1. **Well-calibrated p-values**: As said above, it is possible to proceed with BH procedure. However, BH is known to be slightly overly conservative. Thus, in case of very good to perfect calibration,

it is possible to adjust the BH estimator, so as to limit its conservativeness. This is essentially achieved by adjusting the so-called $\pi_0$ estimate parameter, which quantifies the proportion of null tests (i.e. the proportion of mismatches in the considered set of PSMs), as proposed in [4, 21] in the proteomics context. However, many other $\pi_0$ estimators are available in the literature and a handful of them is directly embedded in CP4P, so that they can be directly used.

2. **Ill-calibrated p-values**: This category contains the cases of intermediate quality, which are the most frequent. Fortunately, in most of the case, the small BH overconservativeness is sufficient to compensate the partial lack of calibration, so that it is nonetheless possible to proceed with the FDR estimate. However, in those cases, ajusting the $\pi_0$ parameter may require an educated eye, so that sticking to the original BH procedure is advised.

3. **Miscalibrated p-values**: In those cases, the lack of calibration is obvious and applying BH procedure will not garantee a trustworthy FDR. However, it is also possible to consider the p-values as classical scores, and to convert them into well-calibrated p-values, following the procedures described above.

Note that a partial lack of calibration may result from the lack of a needed Šidák correction (see Supporting Section S1.2.2); or, on the contrary, from the application of a Šidák correction which was not necessary.

###### S1.2.5 A simple practical tool: CP4P

The principle of Scheweder and Spjøtvoll plot (as well as its CP4P version) is summarized in Figure S3: it is to compute the cumulative sum of 1-($p$-value)s, starting from the largest p-values (i.e. the right hand side of the histogram), so that the horizontal shape of the uniform distribution turns into a slope, which is globally less sensitive to random effects.

With respect to traditional Scheweder and Spjøtvoll plot, the calibration plot proposed in CP4P display visual colored indicator that facilitates its interpretation: In red appears any deviation from the uniform distribution; In green, one indicates the extent to which the p-values of the discoveries (*i.e.* correct matches) are concentrated toward small values; and finally, blue indications refer to the $\pi_0$ parameter, which fine-tuning can help reduce the over-conservativeness of BH original estimator (as illustration, Supporting Section S2 contains many calibration plots resulting from the preliminary checks of p-value calibration, when constructing Figure 1 in the main article).

For more details on CP4P, as well as for tutorials and guidelines on how to use it, we refer previous article of our team: [1] as well as the tutorial available in its supplemental materials (also available there: `https://sites.google.com/site/thomasburgerswebpage/download/tutorial-CP4P-4.pdf`); [10]; and finally, section 4, 5 and 6 of [22], as well as the corresponding supplemental materials (also available there: `https://sites.google.com/site/thomasburgerswebpage/download/suppmat-5tips.pdf`).

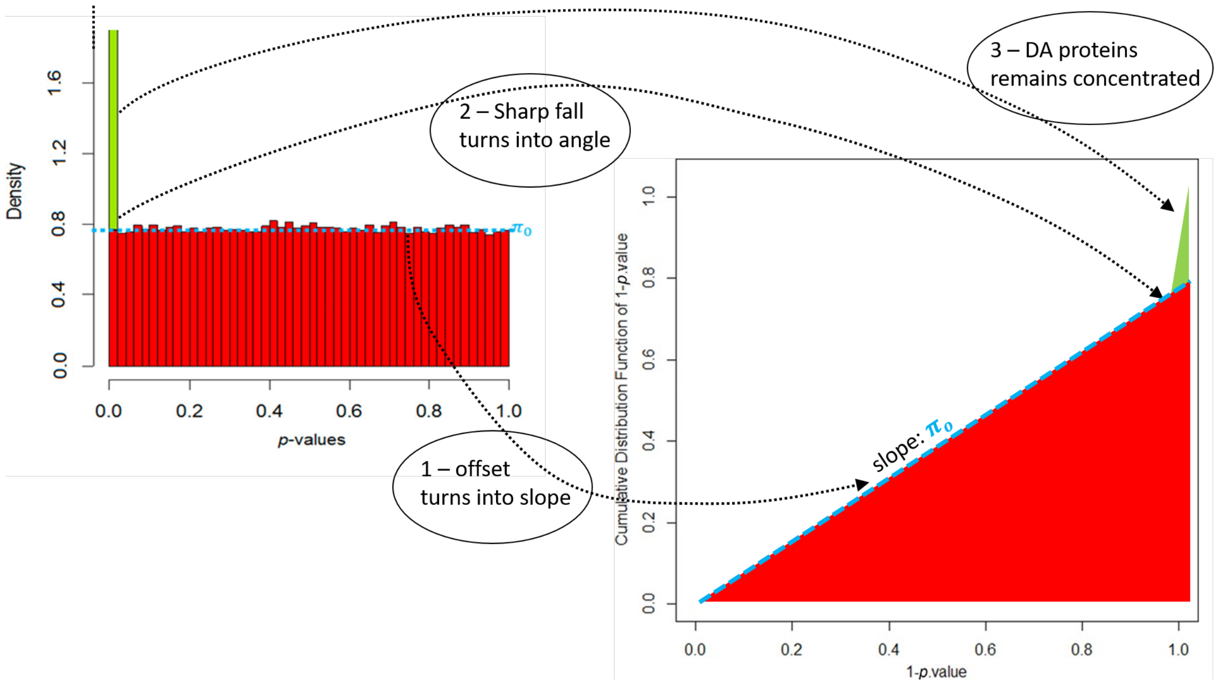

Figure S3: Schematic representation of the calibration plot construction process, starting from the p-value histogram – borrowed from [22].

### S2 Supporting results

#### S2.1 Mascot calibration

This section contains several calibration plots resulting from CP4P (see below, Figures S4 to S15. This package allows adjusting the $\pi_0$ estimate to limit BH over-conservativeness. However, for sakes of general purpose use, we did not rely on this option: We practically forced $\pi_0$ to 1, so as to recover BH original estimator. The plots below correspond to 3 samples (out of the 10 involved in the study): as they are all similar (beyond human eye capabilities), we did not plot them all. We focused on the first one (as the one opening the series), the fourth one (displaying a slightly outlying behavior, see Figure 3 in the main article) and the seventh one (randomly chosen).

The conclusions are as follows: In settings HH, HL and LH the calibrations are excellent: one observes a well-defined elbow, made of a linear part on the left hand side and a sharp increment on the right hand side. However, in the LL setting, the calibration is not as good, without being too bad (notably, no uniformity underestimation appears): This is a small ill-calibration case. However, it contrasts with the 3 other settings. Concretely, it remains possible to compute a BH FDR and to safely rely on it. However, it must remain overly conservative to cope for the approximate calibration. This notably explains why, on Figure 1 (main article), the cut-off scores are higher in the LL setting than in others.

Although not a problem with regards to our investigations and goals, we wondered on this ill-calibration. The point was only to better understand it, not to correct for it (as modifying the p-values in the LL setting would amount to change the Mascot score values, and thus, plotting Figure 1A in the main article would not be possible anymore, as the scales would become different for each setting). However, following Supporting Section S1.2.2, we tried to introduced an additional Šidák correction (as in LL, the number of candidate sequences is higher), but this clearly deteriorated the calibration (see Figure S16). This confirmed us that Mascot is indeed calibrated according to the best-PSM assumption. However, the LL setting being clearly outdated, this is not an issue for Mascot everyday use.

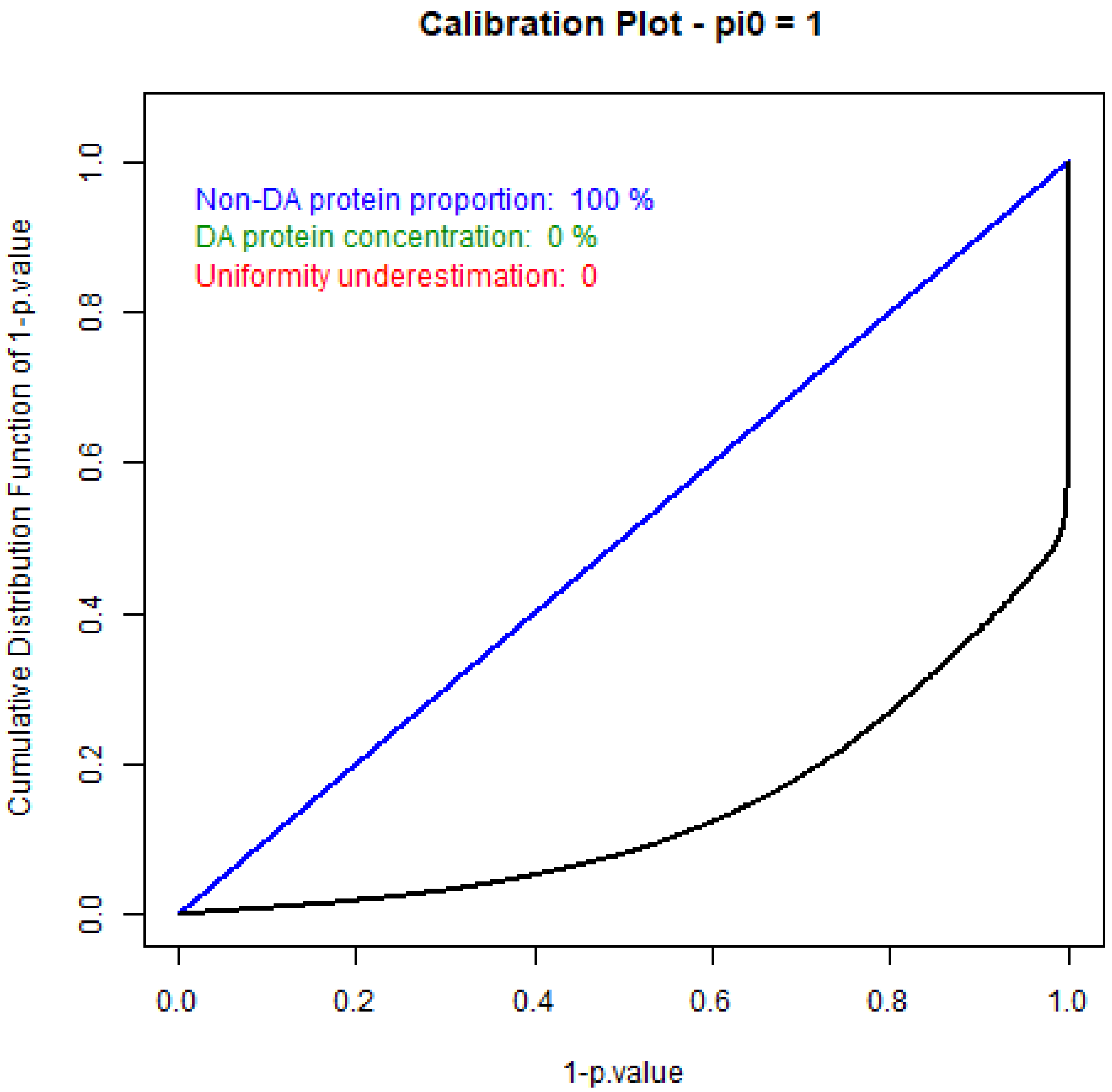

Figure S4: Calibration plot of sample R1 in the LL setting

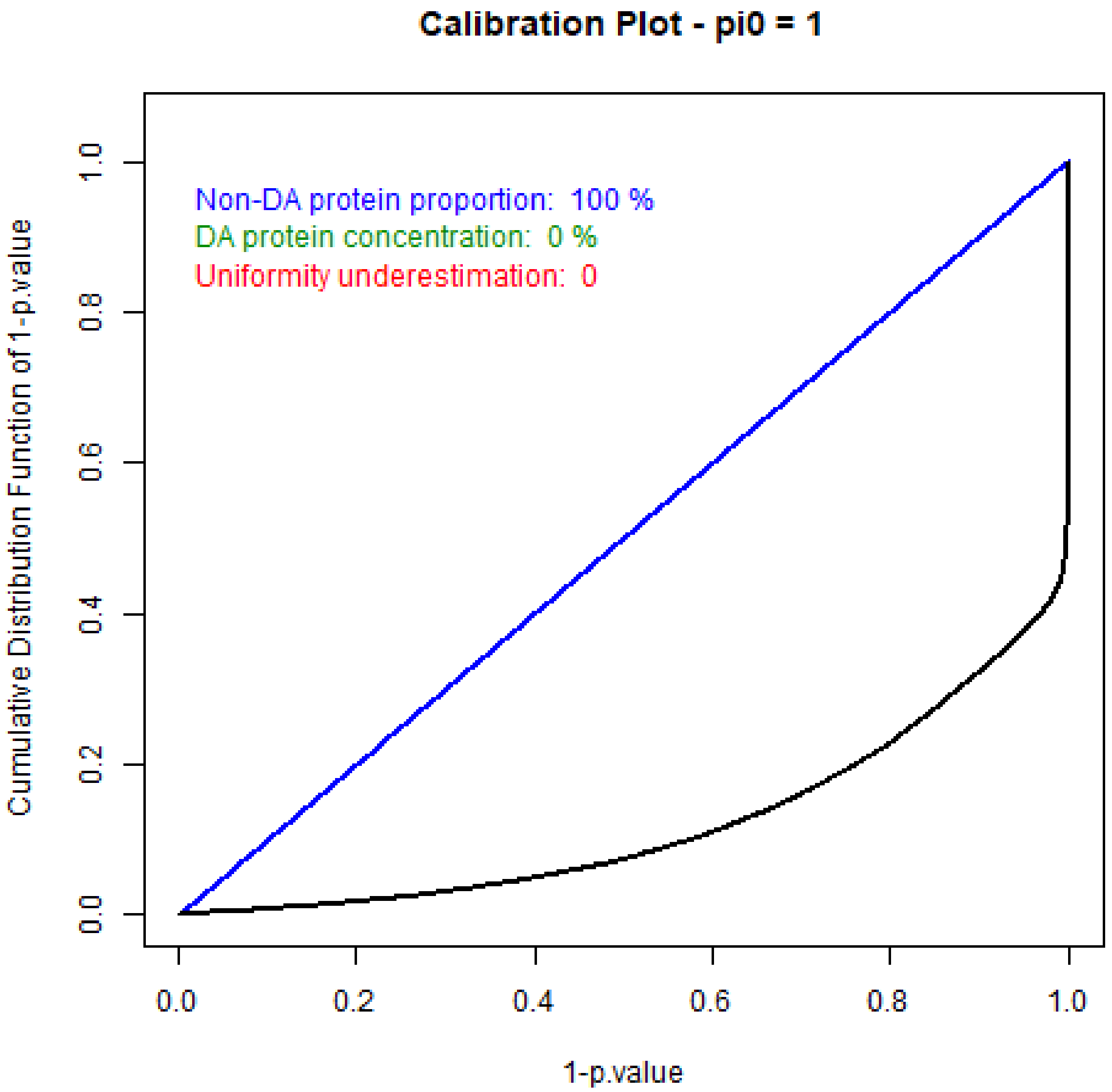

Figure S5: Calibration plot of sample R4 in the LL setting

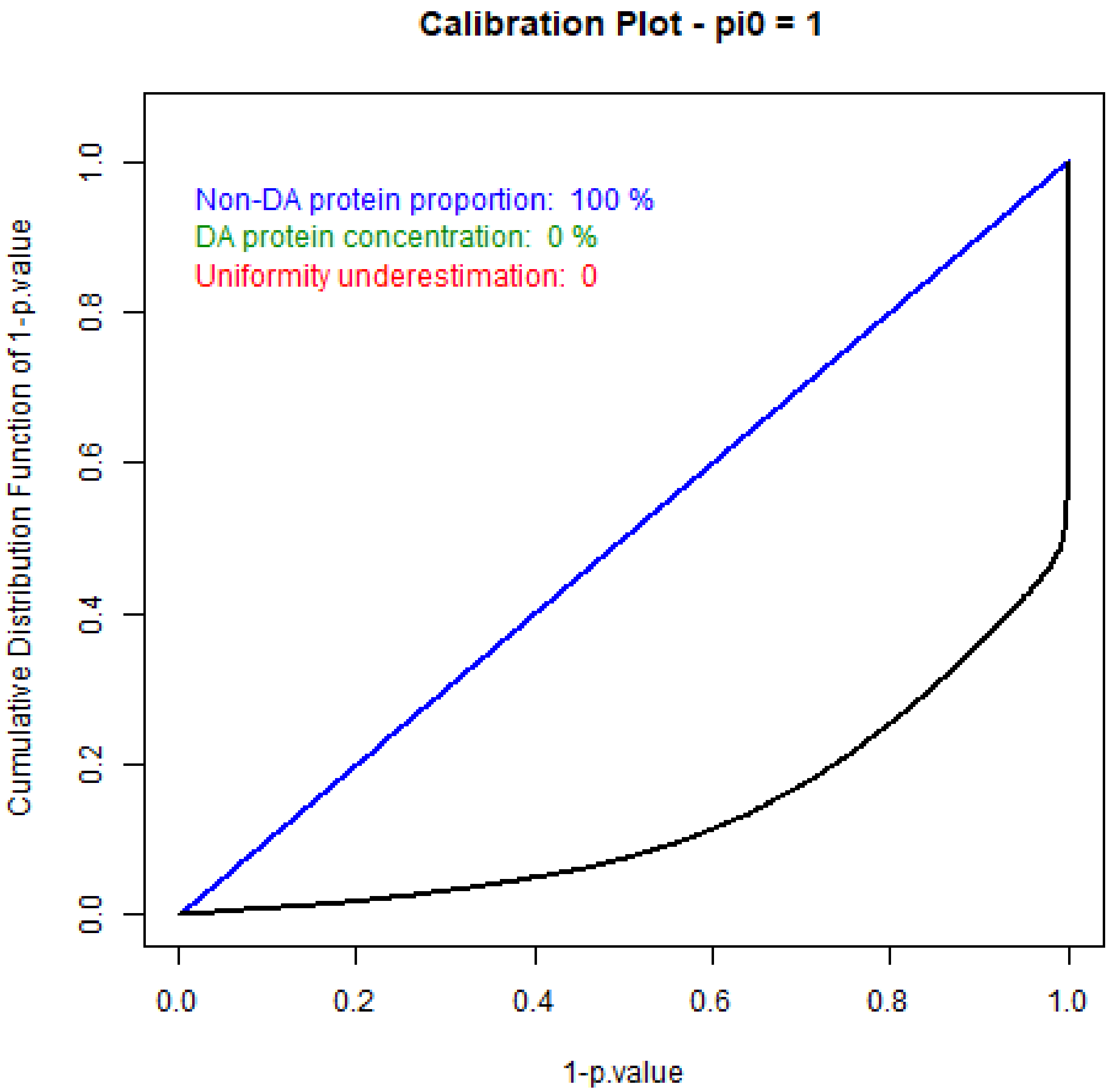

Figure S6: Calibration plot of sample R7 in the LL setting

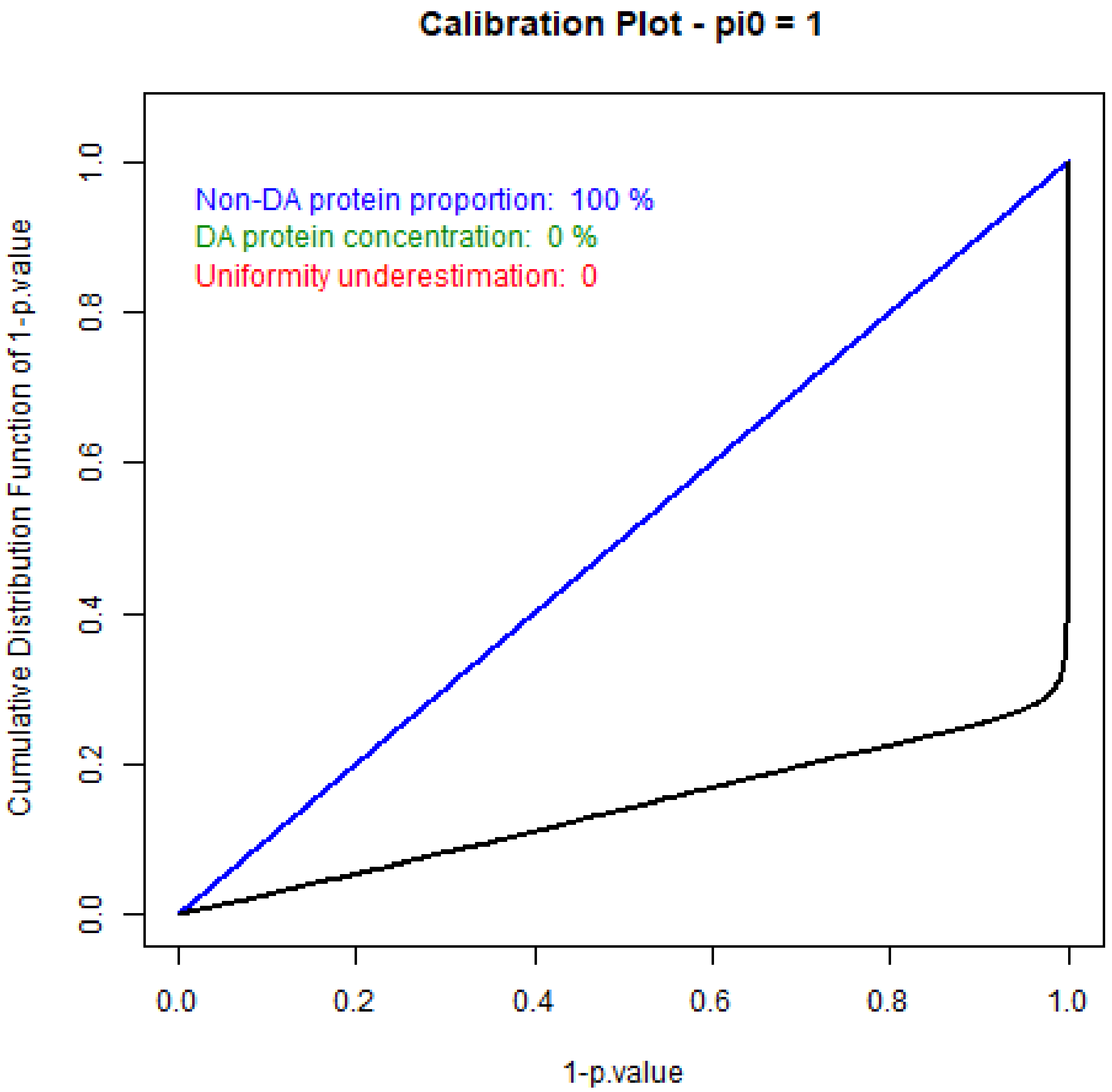

Figure S7: Calibration plot of sample R1 in the HL setting

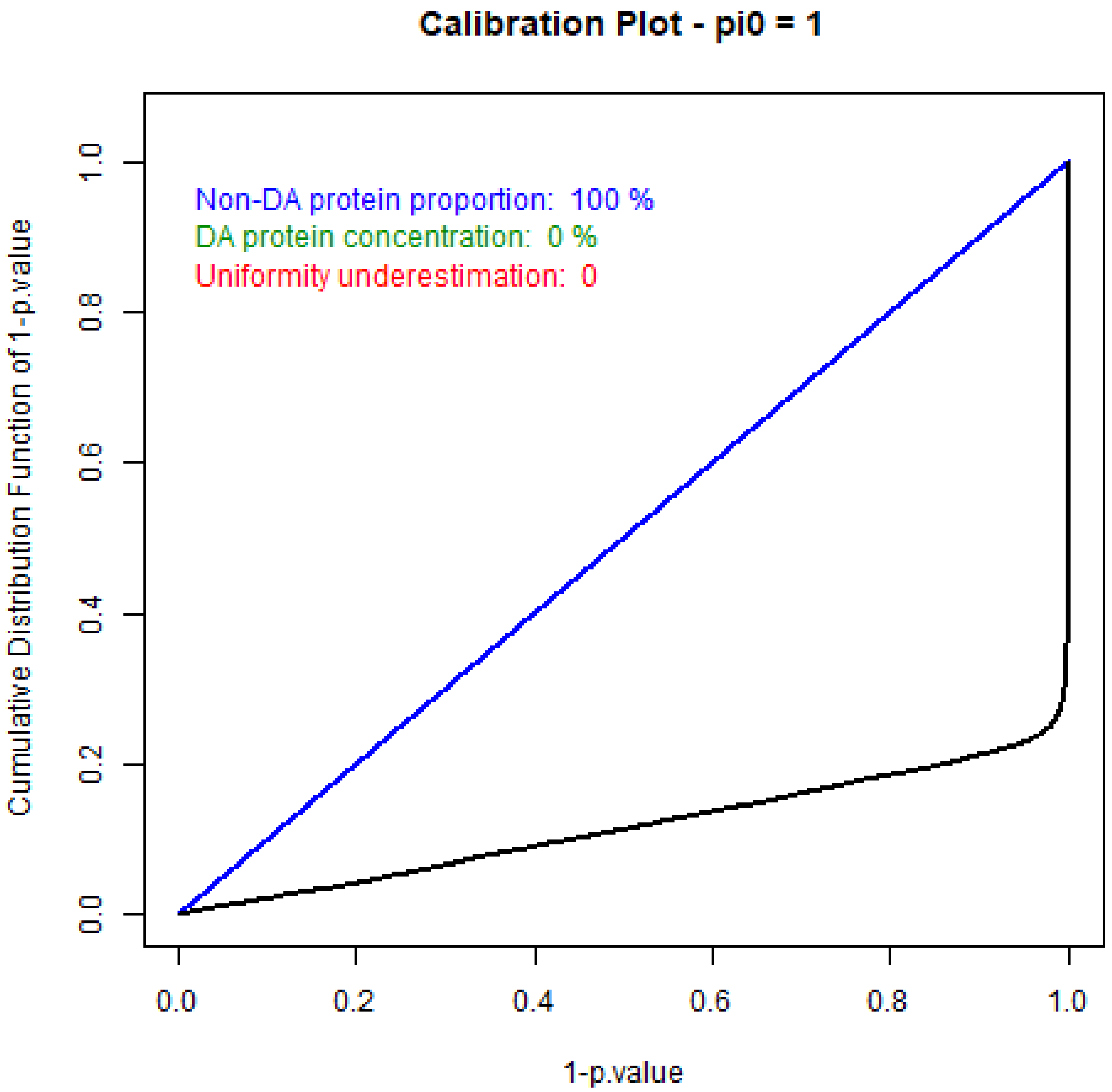

Figure S8: Calibration plot of sample R4 in the HL setting

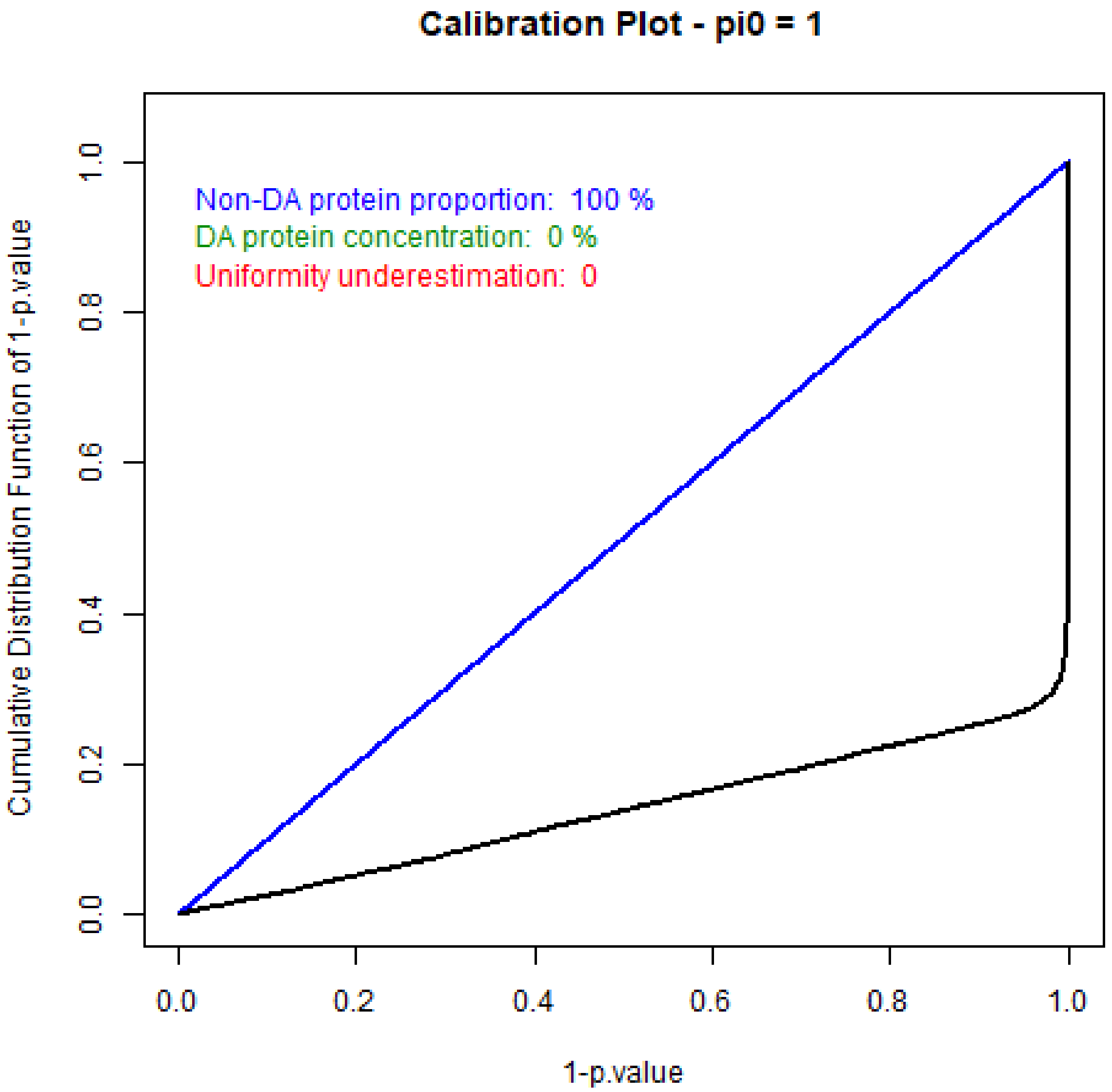

Figure S9: Calibration plot of sample R7 in the HL setting

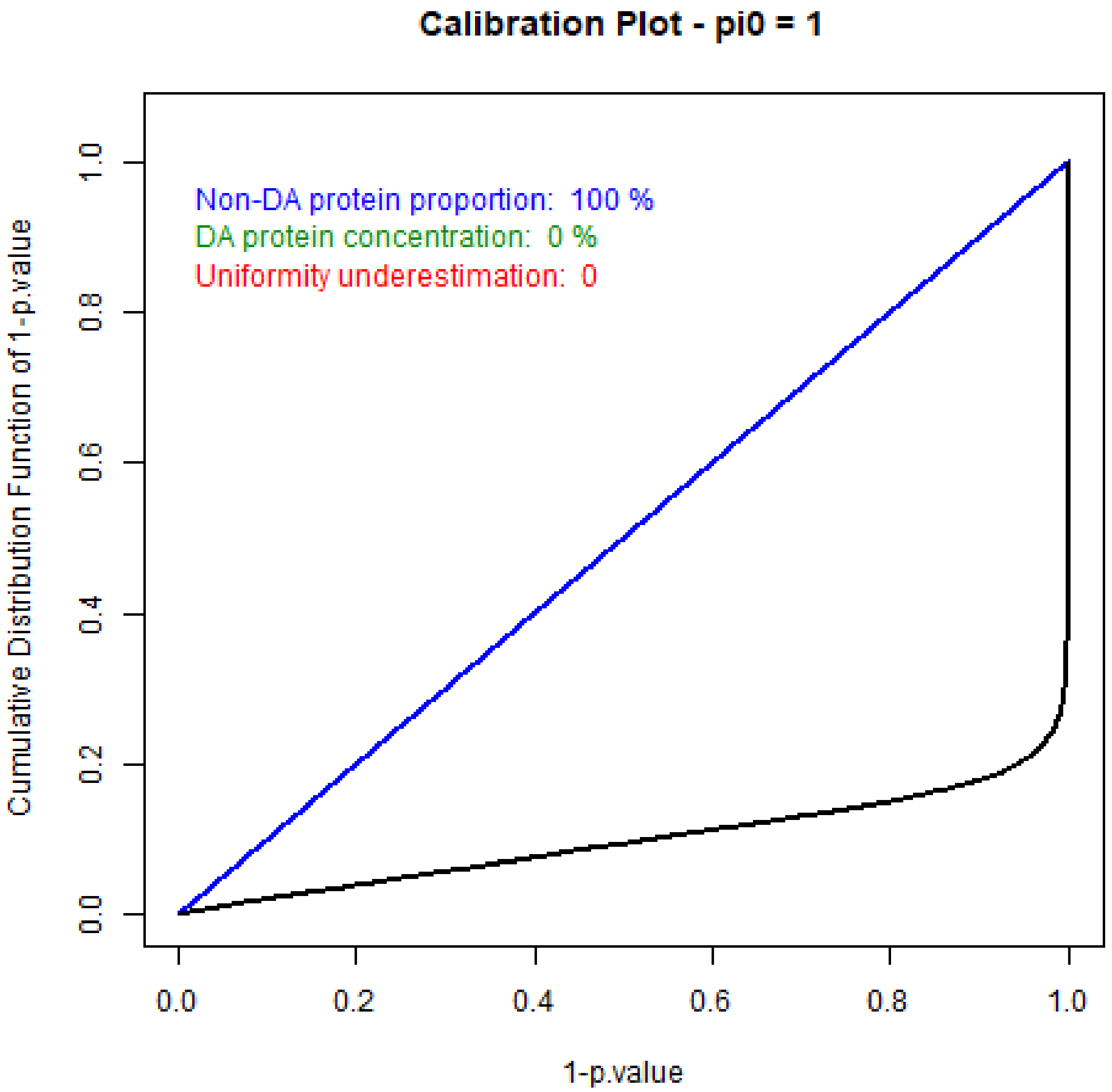

Figure S10: Calibration plot of sample R1 in the LH setting

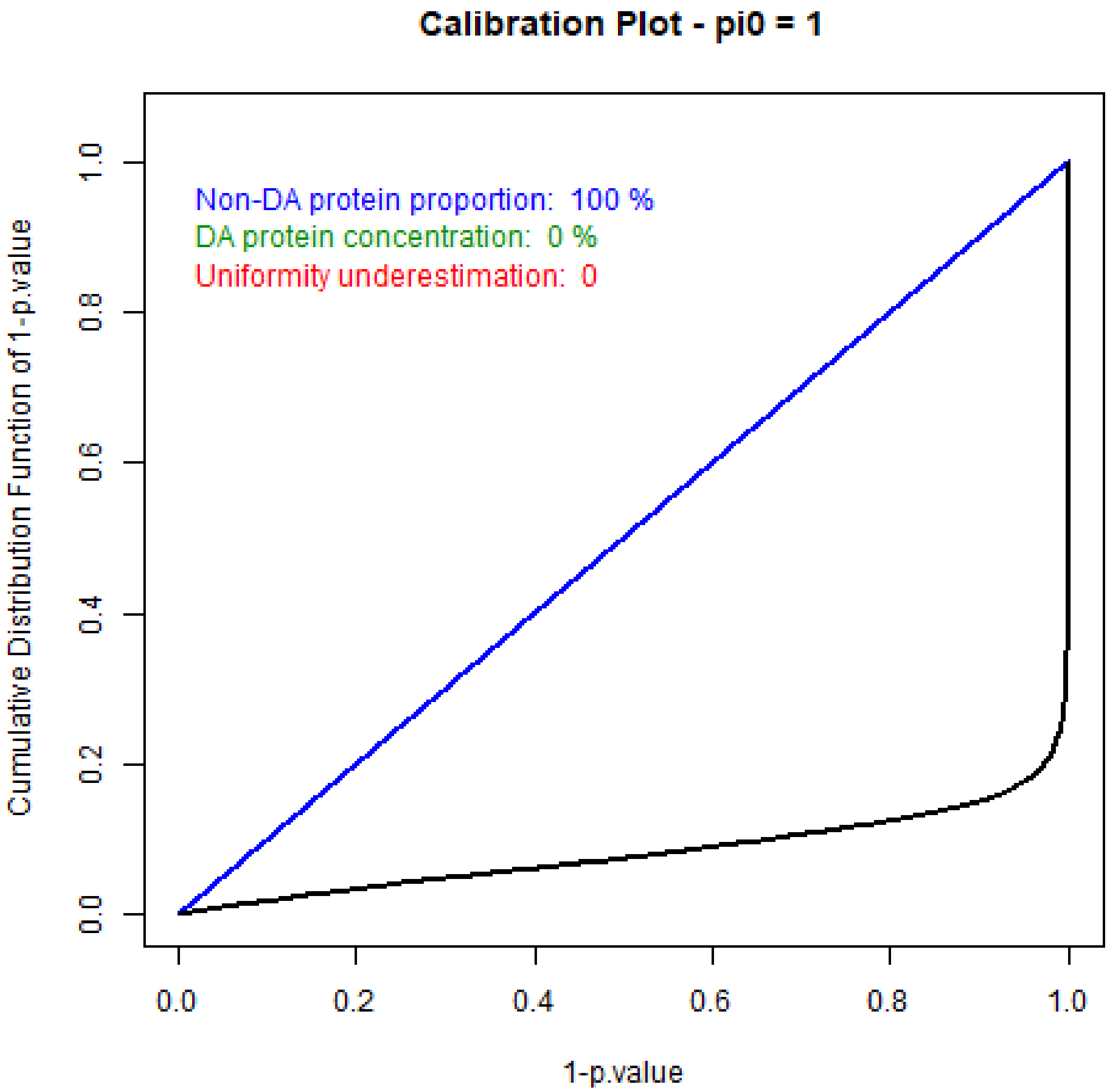

Figure S11: Calibration plot of sample R4 in the LH setting

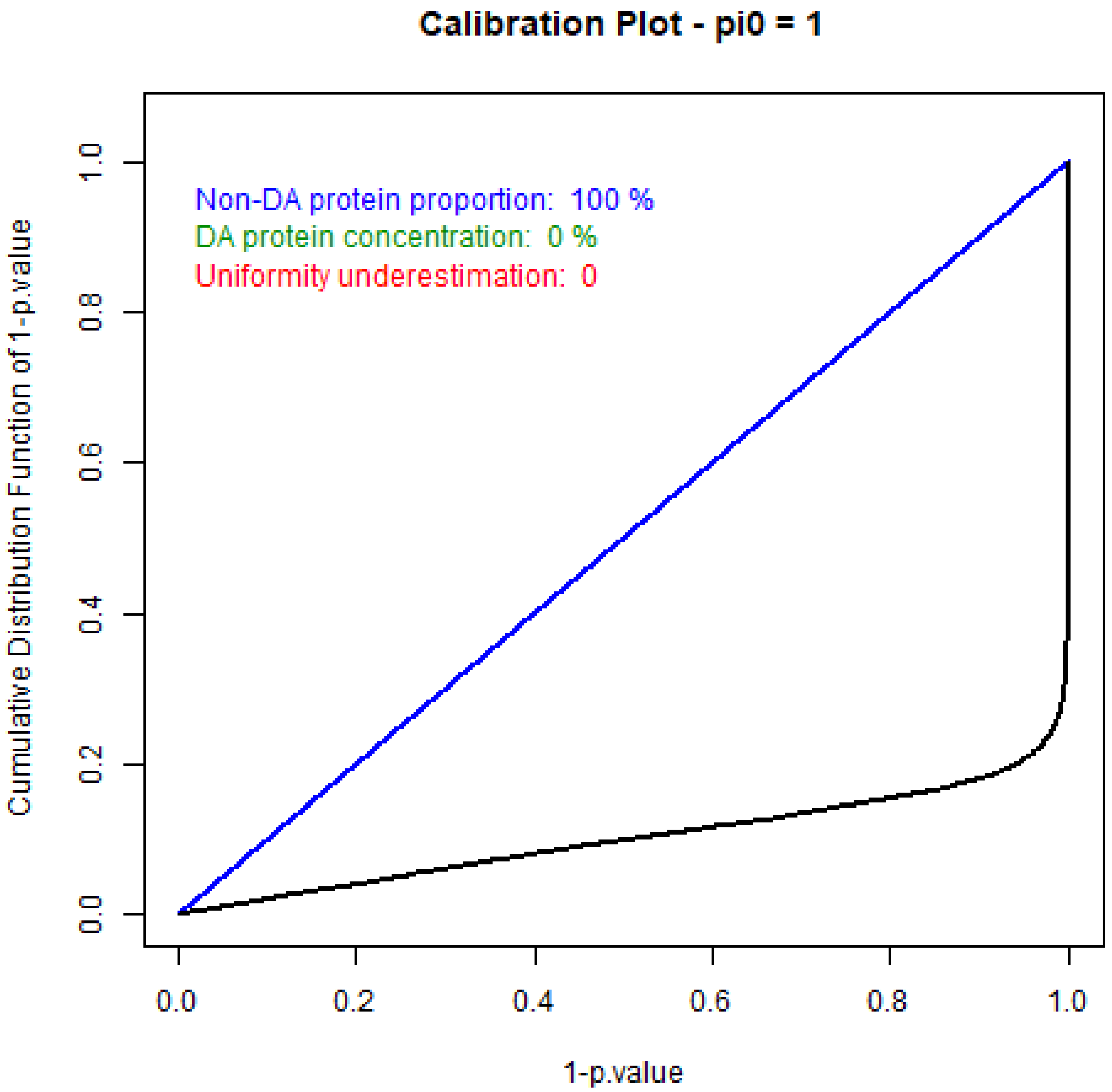

Figure S12: Calibration plot of sample R7 in the LH setting

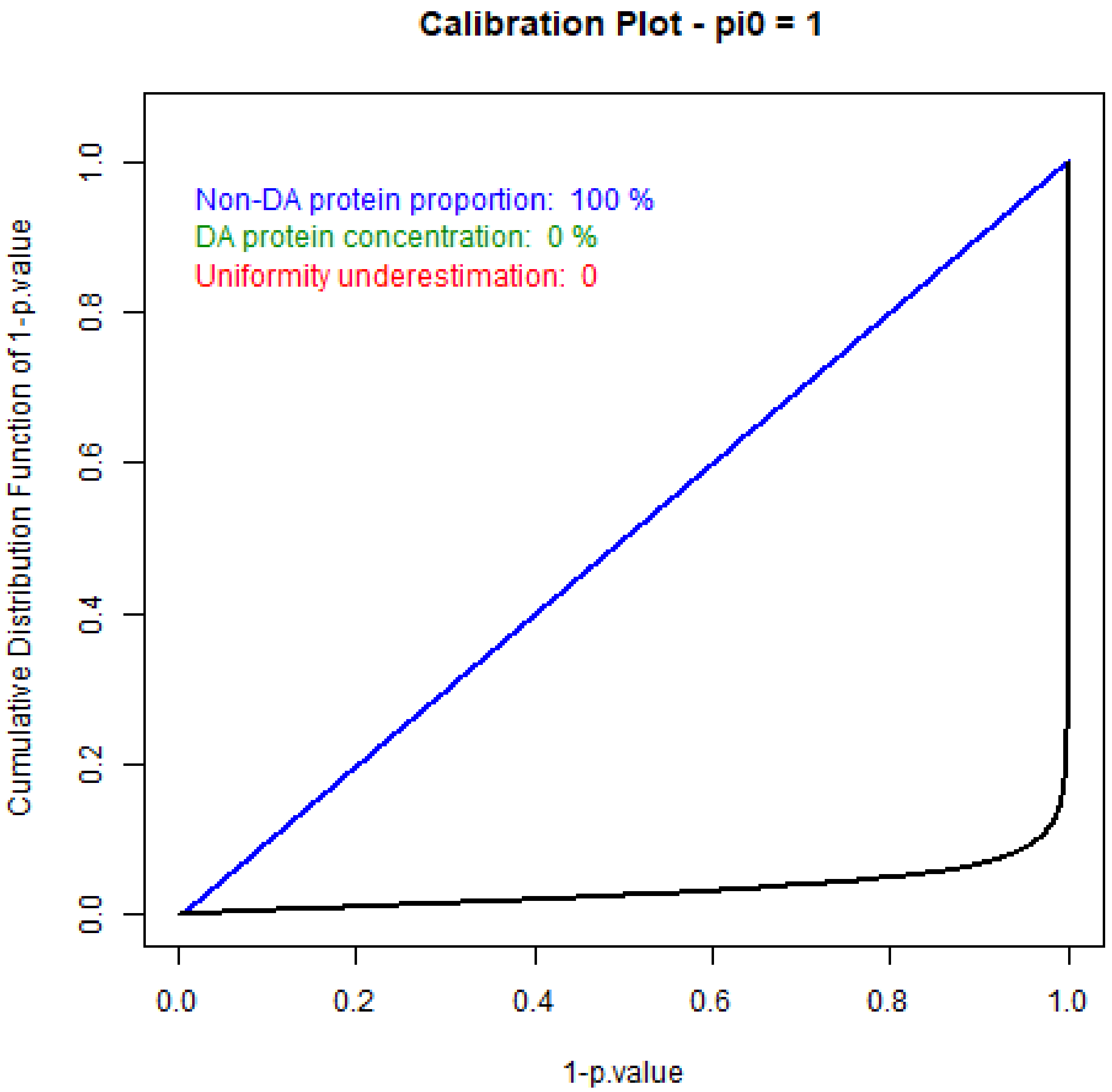

Figure S13: Calibration plot of sample R1 in the HH setting

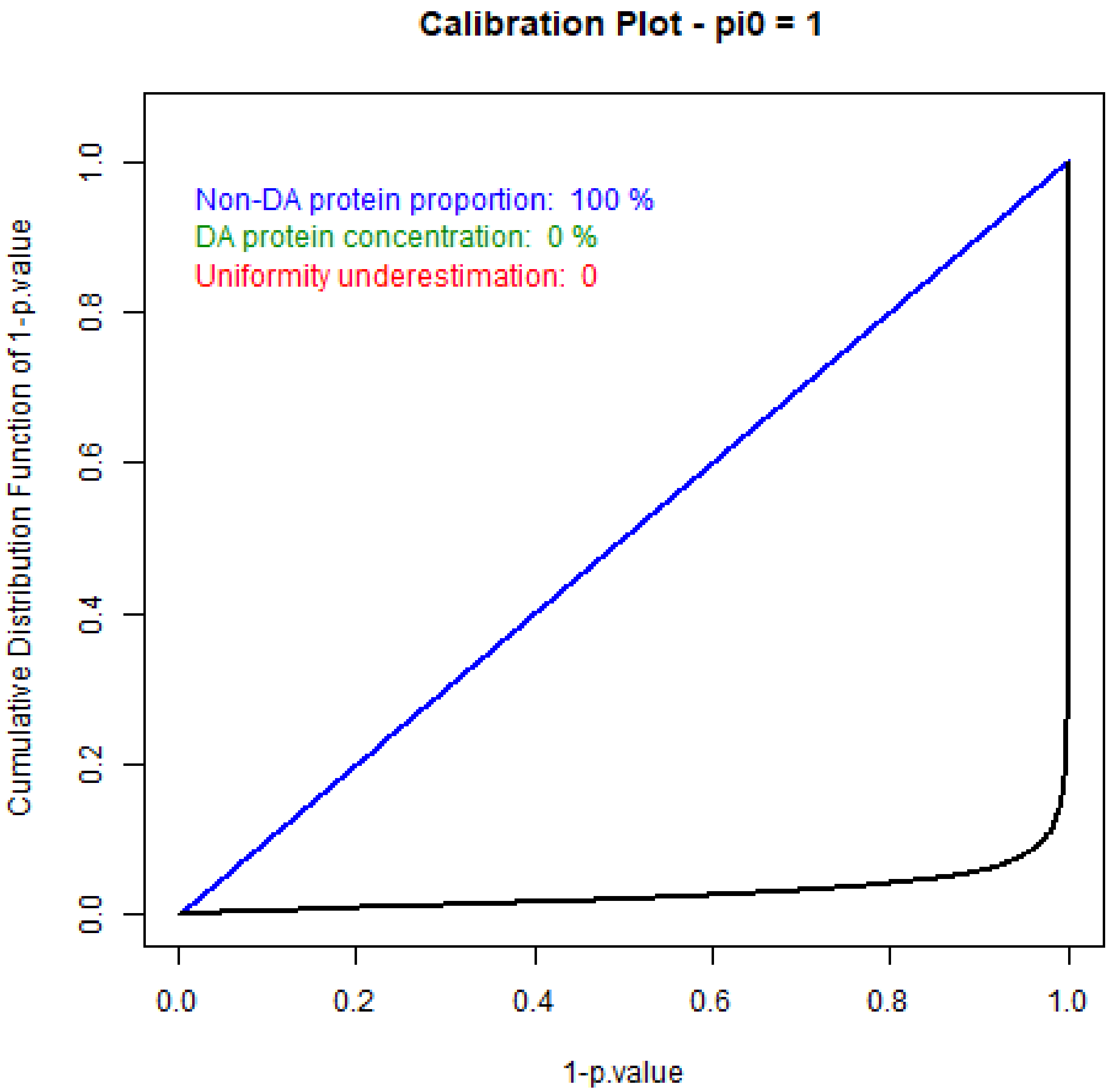

Figure S14: Calibration plot of sample R4 in the HH setting

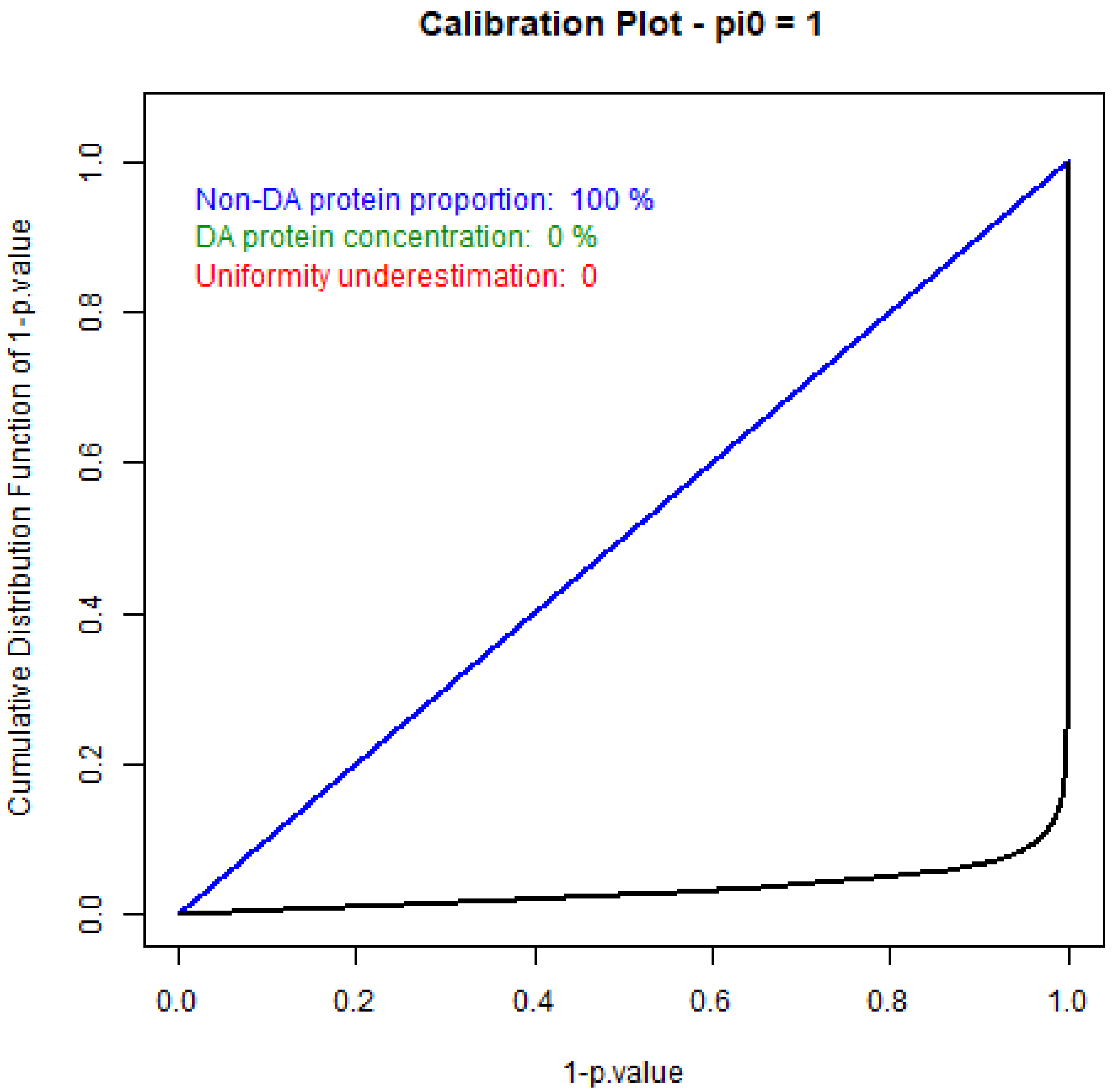

Figure S15: Calibration plot of sample R7 in the HH setting

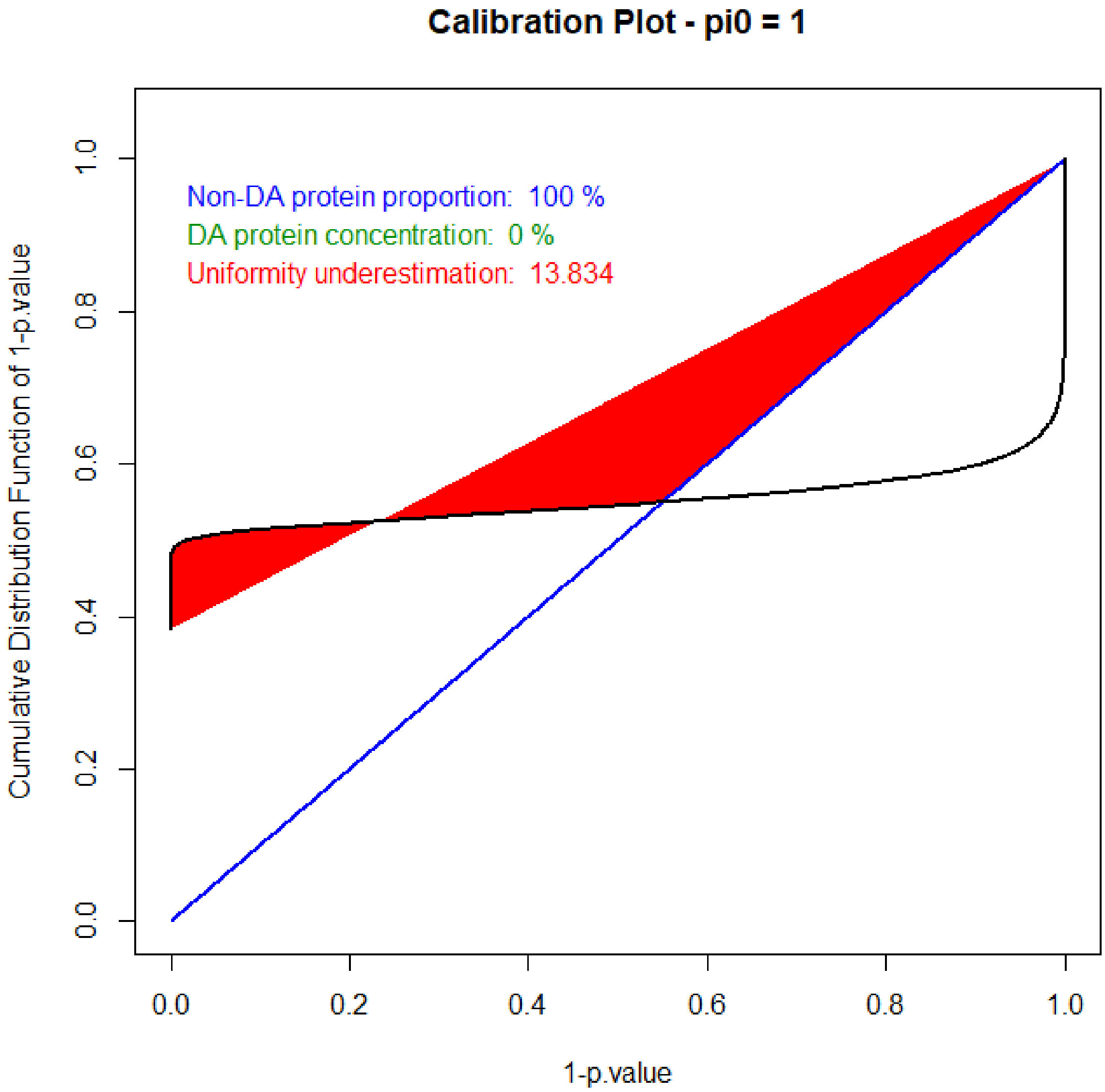

Figure S16: Šidák correction leads to miscalibrated p-values (LL setting, sample R1).

#### S2.2 Miscelaneous supporting tables

Table S1: Quantitative summary of the results depicted in Figure 1, completed with the number of validated PSMs.

| | | | BH | | | | | | | | TDC | | | | | | | |
|---|---|---|---|---|---|---|---|---|---|---|---|---|---|---|---|---|---|---|
| | | | Low-Low | | High-Low | | Low-High | | High-High | | Low-Low | | High-Low | | Low-High | | High-High | |
| | | Queries | PSM target | Score | PSM target | Score | PSM target | Score | PSM target | Score | PSM target | Score | PSM target | Score | PSM target | Score | PSM target | Score |
| QEx | R1 | 26604 | 12135 | 23.21 | 10954 | 21.69 | 11623 | 21.42 | 10551 | 20.71 | 12758 | 19.23 | 11951 | 11.01 | 13363 | 8.83 | 12209 | 1.61 |
| | R2 | 26520 | 11880 | 23.28 | 10863 | 21.78 | 11368 | 21.52 | 10446 | 20.77 | 12546 | 19.41 | 11948 | 11.04 | 13251 | 8.59 | 12224 | 1.58 |
| | R3 | 26103 | 12364 | 23.07 | 10830 | 21.74 | 11863 | 21.41 | 10463 | 20.71 | 12988 | 19.15 | 11802 | 11.04 | 13629 | 8.48 | 12021 | 1.52 |
| | R4 | 24542 | 12680 | 22.68 | 11543 | 21.44 | 12029 | 21.28 | 11045 | 20.71 | 13284 | 18.88 | 12589 | 10.62 | 13828 | 8.69 | 12819 | 1.26 |
| | R5 | 25759 | 11756 | 23.24 | 10696 | 21.76 | 11224 | 21.53 | 10275 | 20.75 | 12309 | 19.64 | 11729 | 10.96 | 12977 | 9.01 | 12039 | 1.2 |
| | R6 | 25900 | 12379 | 23.01 | 11043 | 21.66 | 11853 | 21.39 | 10658 | 20.67 | 12994 | 19.09 | 12049 | 10.64 | 13519 | 9.09 | 12309 | 1.11 |
| | R7 | 25579 | 12121 | 23.07 | 10867 | 21.69 | 11592 | 21.47 | 10488 | 20.70 | 12703 | 19.32 | 11941 | 10.37 | 13276 | 9.05 | 12134 | 1.42 |
| | R8 | 25500 | 12095 | 23.05 | 10825 | 21.65 | 11624 | 21.41 | 10468 | 20.67 | 12659 | 19.26 | 11816 | 10.84 | 13274 | 8.91 | 12019 | 1.39 |
| | R9 | 23726 | 10415 | 23.38 | 9509 | 21.89 | 9975 | 21.69 | 9128 | 20.84 | 10852 | 20.73 | 10439 | 11.66 | 11510 | 10.14 | 10705 | 2.37 |
| | R10 | 23819 | 10428 | 23.43 | 9462 | 21.94 | 9968 | 21.71 | 9145 | 20.82 | 10856 | 20.61 | 10389 | 12.21 | 11507 | 10.45 | 10762 | 2.13 |

| | | | BH | | | | | | | | TDC | | | | | | | |
|---|---|---|---|---|---|---|---|---|---|---|---|---|---|---|---|---|---|---|
| | | | Low-Low | | High-Low | | Low-High | | High-High | | Low-Low | | High-Low | | Low-High | | High-High | |
| | | Queries | PSM target | Score | PSM target | Score | PSM target | Score | PSM target | Score | PSM target | Score | PSM target | Score | PSM target | Score | PSM target | Score |
| Velos | R1 | 18637 | 10784 | 22.17 | 10139 | 21.15 | . | . | . | | 11202 | 19.65 | 11169 | 11.32 | . | . | . | |
| | R2 | 18722 | 10835 | 22.18 | 10210 | 21.22 | . | . | . | | 11179 | 19.96 | 11244 | 11.26 | . | . | . | |
| | R3 | 18511 | 10882 | 22.1 | 10167 | 21.15 | . | . | . | | 11226 | 19.48 | 11127 | 11.24 | . | . | . | |
| | R4 | 18242 | 10848 | 22.06 | 10148 | 21.15 | . | . | . | | 11121 | 19.35 | 11206 | 10.52 | . | . | . | |
| | R5 | 18302 | 10715 | 22.13 | 10048 | 21.19 | . | . | . | | 11166 | 19.25 | 11069 | 11.28 | . | . | . | |
| | R6 | 18352 | 10746 | 22.13 | 10021 | 21.2 | . | . | . | | 11182 | 19.54 | 11084 | 10.93 | . | . | . | |
| | R7 | 18402 | 10794 | 22.13 | 10086 | 21.21 | . | . | . | | 10944 | 20.23 | 11149 | 10.98 | . | . | . | |
| | R8 | 17799 | 10665 | 22.04 | 10008 | 21.17 | . | . | . | | 11725 | 20.91 | 11020 | 10.58 | . | . | . | |
| | R9 | 19041 | 11539 | 22.05 | 10727 | 21.17 | . | . | . | | 11790 | 20.07 | 11658 | 11.96 | . | . | . | |
| | R10 | 19035 | 11499 | 22 | 10676 | 21.15 | . | . | . | | 11084 | 20.22 | 11619 | 11.83 | . | . | . | |

Table S2: Replicate-wise details of the validation results summarized in Table 1.

| | | No validation rule | | | | | | | | |
|---|---|---|---|---|---|---|---|---|---|---|
| | | PSMs | | | Peptides | | | Proteins | | |
| | | #Targets | #Decoys | min Score | #Targets | #Decoys | min Score | #Targets | #Decoys | min Score |
| Fivefold shuffle | R1 | 12319 | 744 | 0 | 9293 | 734 | 0 | 1482 | 712 | 0 |
| | R2 | 12347 | 840 | 0 | 9286 | 821 | 0 | 1462 | 795 | 0 |
| | R3 | 12149 | 849 | 0.01 | 9254 | 829 | 0.01 | 1451 | 803 | 0.01 |
| | R4 | 12843 | 714 | 0.01 | 9898 | 684 | 0.01 | 1427 | 667 | 0.01 |
| | R5 | 12092 | 826 | 0 | 9465 | 806 | 0 | 1470 | 775 | 0 |
| | R6 | 12332 | 837 | 0 | 9422 | 821 | 0 | 1435 | 784 | 0 |
| | R7 | 12201 | 852 | 0 | 9408 | 833 | 0 | 1476 | 807 | 0 |
| | R8 | 12095 | 830 | 0 | 9418 | 808 | 0 | 1477 | 780 | 0 |
| | R9 | 10908 | 935 | 0 | 9026 | 914 | 0 | 1503 | 869 | 0 |
| | R10 | 10916 | 946 | 0 | 9043 | 922 | 0 | 1482 | 868 | 0 |
| | Average | 12020.2 | 837.3 | 0.002 | 9351.3 | 817.2 | 0.002 | 1466.5 | 786 | 0.002 |

| | | 1% FDR at PSM level (alone) | | | | | | | | |
|---|---|---|---|---|---|---|---|---|---|---|
| | | PSMs | | | Peptides | | | Proteins | | |
| | | #Targets | #Decoys | min Score | #Targets | #Decoys | min Score | #Targets | #Decoys | min Score |
| Fivefold shuffle | R1 | 10523 | 9 | 20.94 | 8175 | 9 | 18.9 | 1305 | 9 | 21.04 |
| | R2 | 10420 | 8 | 21.02 | 8115 | 8 | 20.17 | 1293 | 8 | 21.78 |
| | R3 | 10424 | 11 | 20.96 | 8151 | 11 | 19.84 | 1286 | 11 | 20.96 |
| | R4 | 11012 | 29 | 20.9 | 8723 | 23 | 19.61 | 1292 | 22 | 21.01 |
| | R5 | 10239 | 11 | 21.02 | 8250 | 11 | 19.96 | 1304 | 11 | 21.23 |
| | R6 | 10624 | 10 | 20.93 | 8309 | 10 | 19.55 | 1288 | 10 | 20.95 |
| | R7 | 10458 | 10 | 20.98 | 8272 | 10 | 20.23 | 1305 | 9 | 21.08 |
| | R8 | 10440 | 10 | 20.95 | 8327 | 10 | 19.73 | 1318 | 10 | 20.95 |
| | R9 | 9094 | 9 | 21.15 | 7723 | 9 | 18.53 | 1290 | 9 | 21.54 |
| | R10 | 9102 | 8 | 21.15 | 7758 | 8 | 20.24 | 1296 | 8 | 21.33 |
| | Average | 10233.6 | 11.5 | 21 | 8180.3 | 10.9 | 19.676 | 1297.7 | 10.7 | 21.187 |

| | | 1% FDR at peptide level (alone) | | | | | | | | |
|---|---|---|---|---|---|---|---|---|---|---|
| | | PSMs | | | Peptides | | | Proteins | | |
| | | #Targets | #Decoys | min Score | #Targets | #Decoys | min Score | #Targets | #Decoys | min Score |
| Fivefold shuffle | R1 | 11073 | 8 | 0.04 | 8151 | 8 | 20.92 | 1304 | 8 | 21.04 |
| | R2 | 11032 | 8 | 0.02 | 8089 | 8 | 20.97 | 1293 | 8 | 21.78 |
| | R3 | 10926 | 11 | 0.02 | 8126 | 11 | 20.95 | 1286 | 11 | 20.96 |
| | R4 | 11537 | 32 | 0.02 | 8705 | 23 | 20.85 | 1292 | 22 | 21.01 |
| | R5 | 10755 | 10 | 0 | 8232 | 10 | 20.96 | 1304 | 10 | 21.23 |
| | R6 | 11117 | 9 | 0.01 | 8286 | 9 | 20.92 | 1287 | 9 | 20.95 |
| | R7 | 10975 | 9 | 0.02 | 8251 | 9 | 20.95 | 1305 | 8 | 21.08 |
| | R8 | 10909 | 9 | 0.01 | 8307 | 9 | 20.9 | 1318 | 9 | 20.95 |
| | R9 | 9484 | 8 | 0.05 | 7698 | 8 | 21.12 | 1290 | 8 | 21.54 |
| | R10 | 9554 | 7 | 0.04 | 7750 | 7 | 21.09 | 1300 | 7 | 21.09 |
| | Average | 10736.2 | 11.1 | 0.023 | 8159.5 | 10.2 | 20.963 | 1297.9 | 10 | 21.163 |

| | | 1% FDR at protein level (alone) | | | | | | | | |
|---|---|---|---|---|---|---|---|---|---|---|
| | | PSMs | | | Peptides | | | Proteins | | |
| | | #Targets | #Decoys | min Score | #Targets | #Decoys | min Score | #Targets | #Decoys | min Score |
| Fivefold shuffle | R1 | 12123 | 7 | 0.04 | 9099 | 7 | 0.08 | 1300 | 7 | 22.54 |
| | R2 | 12163 | 8 | 0.02 | 9103 | 8 | 0.03 | 1293 | 7 | 22.7 |
| | R3 | 11964 | 14 | 0.01 | 9070 | 14 | 0.01 | 1282 | 11 | 22.45 |
| | R4 | 12702 | 32 | 0.02 | 9760 | 21 | 0.04 | 1293 | 18 | 22.25 |
| | R5 | 11912 | 13 | 0 | 9288 | 12 | 0.01 | 1299 | 9 | 22.35 |
| | R6 | 12170 | 16 | 0.01 | 9259 | 14 | 0.01 | 1284 | 9 | 22.44 |
| | R7 | 12019 | 13 | 0 | 9227 | 12 | 0 | 1306 | 8 | 22.48 |
| | R8 | 11924 | 12 | 0 | 9247 | 11 | 0 | 1316 | 8 | 22.42 |
| | R9 | 10679 | 12 | 0.02 | 8801 | 10 | 0.02 | 1293 | 7 | 22.99 |
| | R10 | 10710 | 11 | 0.01 | 8840 | 11 | 0.01 | 1290 | 8 | 22.72 |
| | Average | 11836.6 | 13.8 | 0.013 | 9169.4 | 12 | 0.021 | 1295.6 | 9.2 | 22.534 |

#### S2.3 Downfall of Andromeda scores

The TDC lack of stability reported with Mascot in the main article (see Figure 1) can also be illustrated with Andromeda, even though the conclusions should be cautiously interpreted for the following reasons: (1) Andromeda code is not accessible, so that it is not possible to check whether the provided scores are individual or if they should be considered as contextualized scores (see Supporting Information S4.1), because of the "*fixed additive component*" which accounts for peptide dependences, as described in [23]; (2) Depending on the version, the TDC procedure is by default applied on posterior probabilities or on delta scores, both of us being contextualized scores; (3) Due to the specific relationship between the posterior probabilities and the scores of long peptides, the minimum observed Andromeda score within the validated list is almost always near zero, so that focusing on the variations of cut-off score is not informative.

Concerning the first point, it is more a state of affair than an issue to solve, which will only make Andromeda more or less adapted to evaluate the TDC procedure. This is notably the reason why we do not compute and display the BH cut-off scores: it is impossible to make sure that Equation 1 (in the article) is applied on the correct score. As for the second one, we have used the posterior probability, because it is the published method. Finally, concerning the last one, we simply have to find a statistics other than the minimum score which depicts the quality of the borderline validated peptides. We have decided to rely on the lowest percentile of the score distribution. This interprets as following: a value of $x$ indicates that the 1% lowest PSM scores are distributed between 0 and $x$. As detailed in the main article, as well as in Supporting Information S4.1, as the lowest PSM scores are expected to remain of constant quality, it makes an interesting statistics to illustrate a potential lack of stability in the TDC.

The results are depicted on Figure S17. Although the first percentile can be expected to be more stable than the minimum value, one observes an important instability, both for each tolerance tuning taken individually, and across the tolerance tunings. However, it appears that contrarily to Mascot, X!tandem and MS-GF+, the effect of the fragment mass tolerance tuning is much more important than that of the precursor one. Moreover, the mapping between the Velos and Qex analyses is not as good as with Mascot. However, in the LL setting, switching from a Velos to a Qex leads to a stringency loss, while on the contrary, it leads to a stringency increment in the HL setting. Considered together, these observations confirms the lack of of stability of TDC procedure.

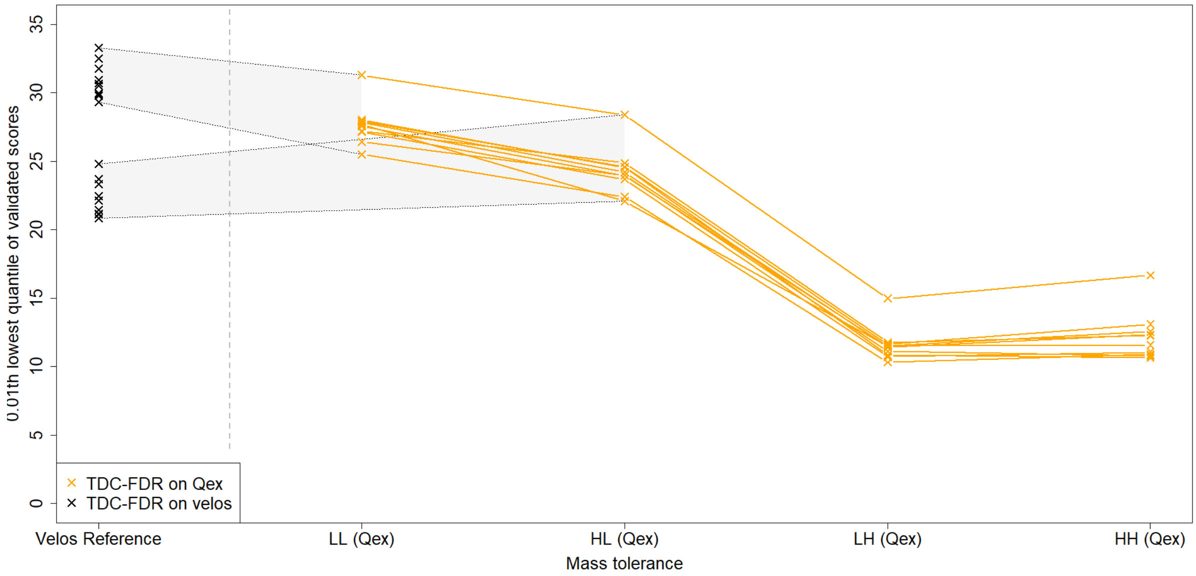

Figure S17: Same figure as Figure 1, yet with Andromeda search engine instead of Mascot.

#### S2.4 TDC can lead to the validation of unreliable PSMs

For illustration purpose, the following figures (Figures S18 to S25) display 8 randomly chosen PSMs with scores lower than 10, out of the 150 ones which are discussed in the "TDC instability can lead to anti-conservative FDRs" section of the article. For sakes of readability, we did not included the 150 PSMs as figures in the present supporting information file. However, they are all stored as PNG files in the companion zipped folder (Supporting Information S5).

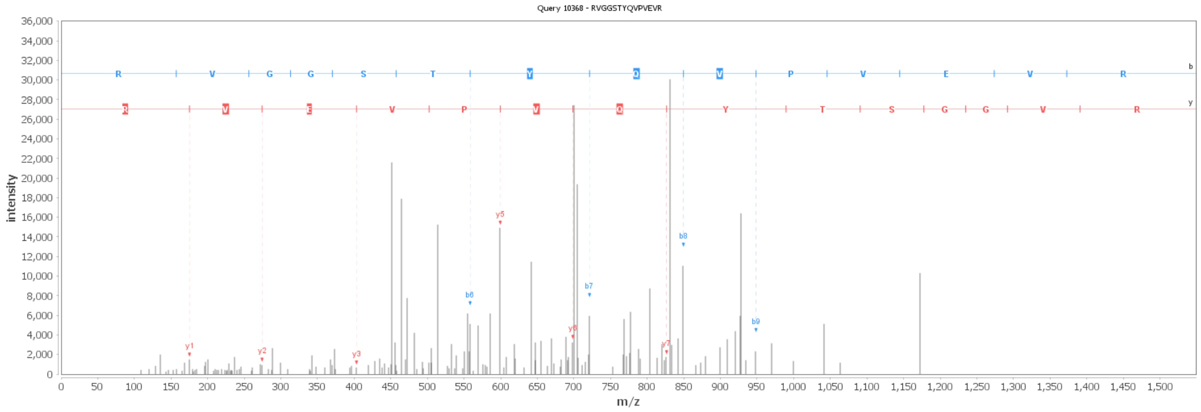

Figure S18: Dubious PSM with score 1.64.

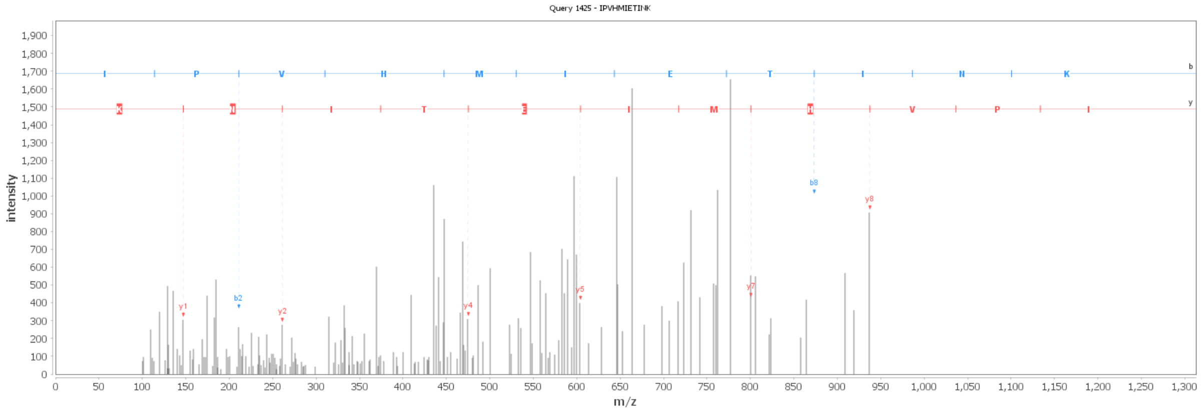

Figure S19: Dubious PSM with score 3.65.

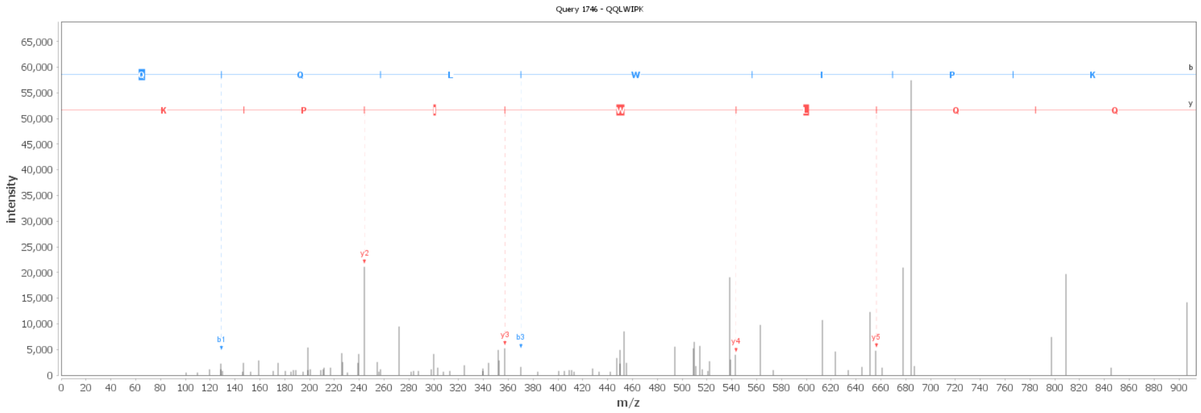

Figure S20: Dubious PSM with score 4.14.

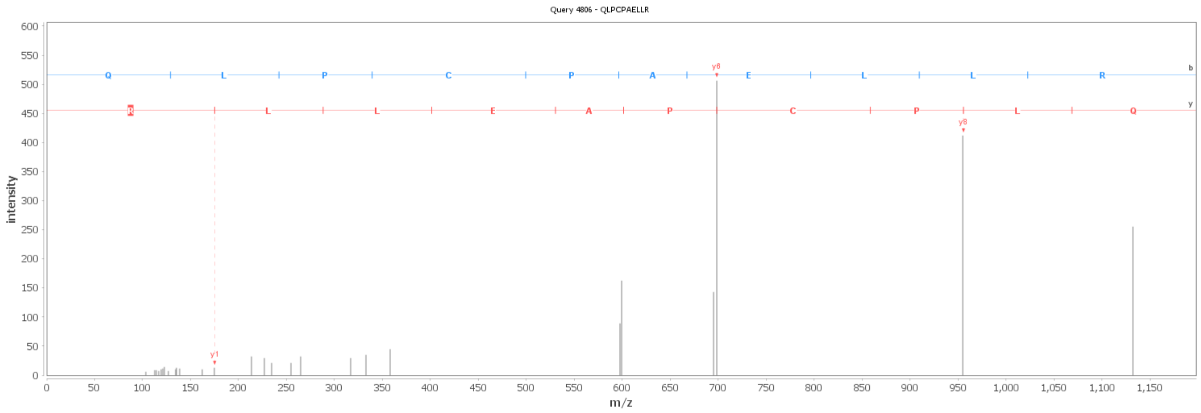

Figure S21: Dubious PSM with score 4.88.

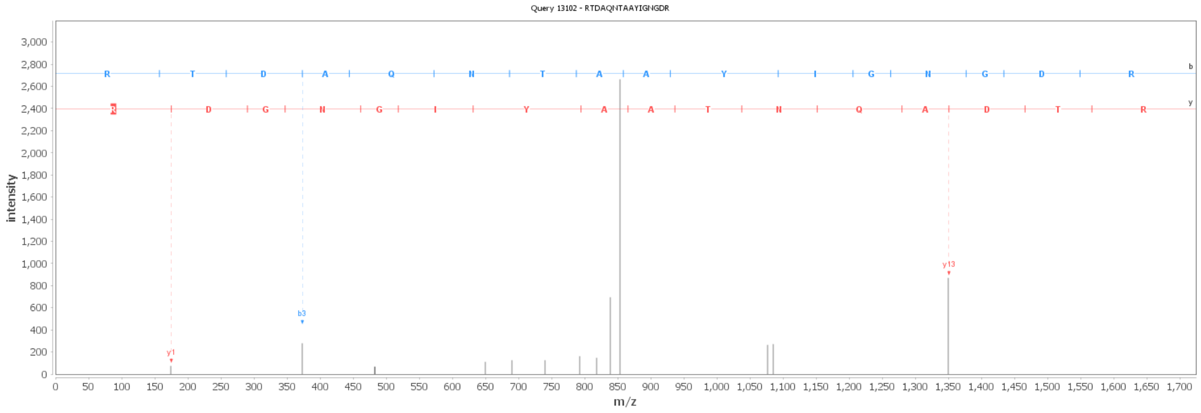

Figure S22: Dubious PSM with score 5.78.

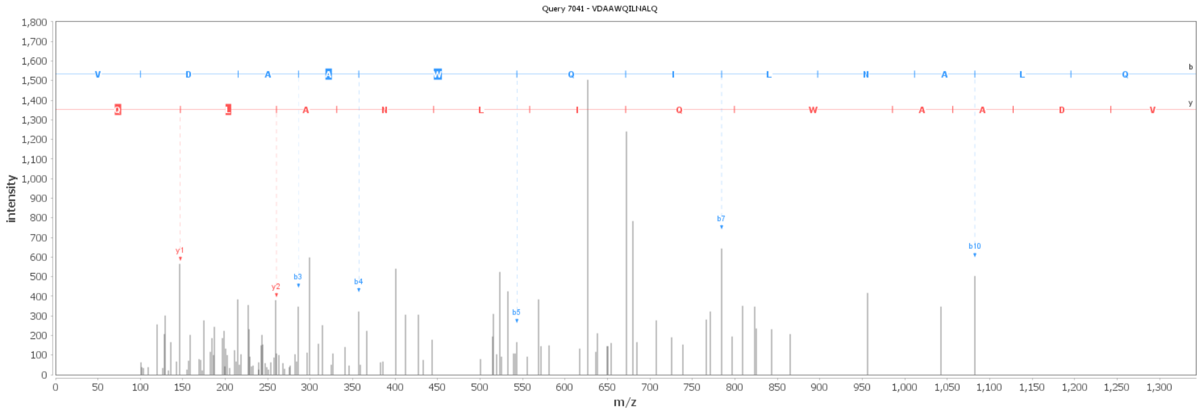

Figure S23: Dubious PSM with score 6.85.

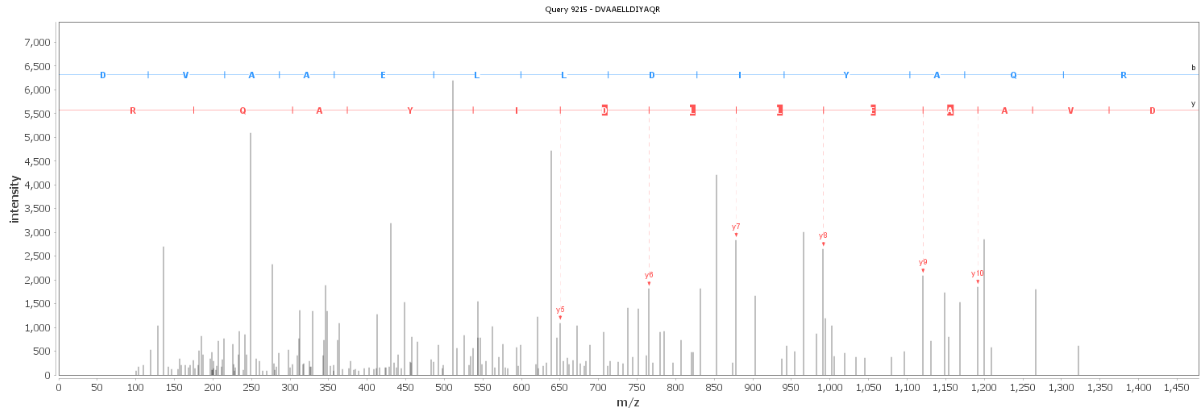

Figure S24: Dubious PSM with score 8.10.

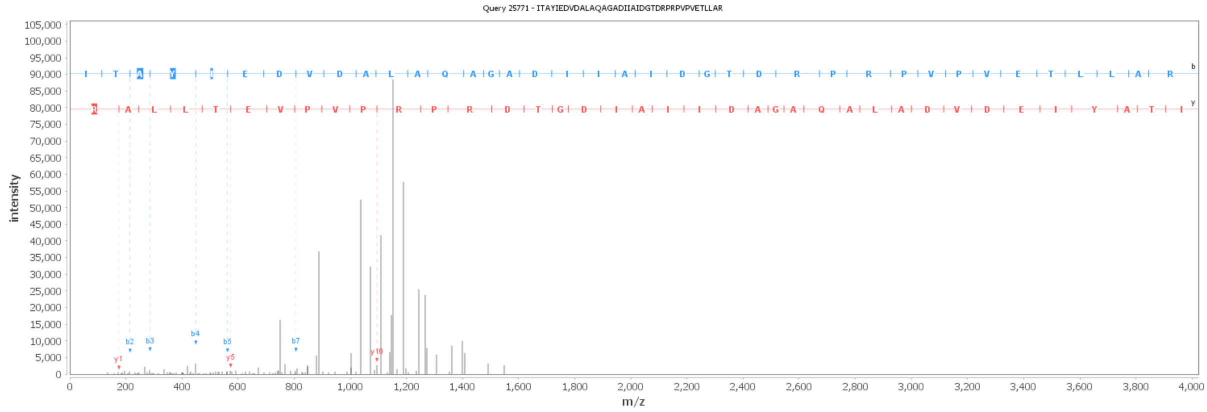

Figure S25: Dubious PSM with score 9.30.

### S3 Supporting methods

#### S3.1 Inference rules

*Peptide inference* or *protein inference* are umbrella terms which encompass several distinct notions: Inference rules, scoring methodologies and quality control metrics. The **inference rules** define which pieces of information should be considered and how they should be used in the inference process, regardless of their quality or reliability. For the spectrum-to-peptide inference, this notably refers to the possibly multiple ranked interpretation of a spectrum. For the peptide-to-protein inference, this refers to the minimum number of peptides per protein, the processing of shared peptides and the definition of protein groups. The **scoring methodology** refers to the definition of a confidence value for each entity (defined thanks to the inference rules), in order to rank them from the most confident to the least one. Finally, the **quality control metrics** is used to filter out some insufficiently reliable entities so as to keep a shorter list of validated ones. The metrics can either be individual (each entity is considered independently of the others) such as with Posterior Error Probability [24, 25]; or associated to the entire filtered list (typically, an FDR, but other multiple test corrections methods exist [26]).

Although conceptually distinct, these notions can overlap in practice, see [27, 15, 16]: Some inference rules directly involve the scoring methodology; Quality control metrics may tightly relate to the scoring methodology; Inference rules and scoring systems are compared so as to find the combination leading to the lowest FDRs; Etc. However, for sake of generality, we kept here a clear distinction. Concretely: *(i)* We did not address the definition of inference rules, and we considered the most simple one (*i.e.* a single peptide interpretation per spectra and only protein-group specific peptides, regardless their number and the protein grouping), and leave to future work (or to any inspired reader) the application of our procedure to more elaborated inference rules; *(ii)* We focused on the scoring methodology, with the objective to preserve the $[0,1]$-uniform distribution, so as to call them well-calibrated peptide/protein p-values; *(iii)*

Regarding the quality control metrics, we obviously relied on BH procedure, which becomes possible thanks to the calibration correctness.

This complete separation provides two advantages. First, it makes it possible to reason on each step independently of the others. Notably, this article focuses on the scoring methodology independently of the inference rules and the quality control metrics. Second, it enables a distinction between the quality control level and and the nature of the controlled entities: While it is customary to validate a list of proteins with an FDR of 1%, it is not as classical to validate a list of PSMs with the criterion that less than 1% of the proteins they map on are putatively false discoveries. However, as illustrated in the Discussion, such options are really insightful.

#### S3.2 Peptide score definition

**Definition 1** *Let us have a peptide sequence* $\mathbf{seq}_i$*, a spectrum* $\mathbf{spec}_j$ *and a score reading*

$$S^{\circ}_{ij} = \mathtt{Score}(\mathbf{seq}_i, \mathbf{spec}_j) \tag{1}$$

*that is provided by a search engine. The triplet* $(\mathbf{seq}_i, \mathbf{spec}_j, S^{\circ}_{ij})$ *formally defines a Peptide-Spectrum-Match (or a PSM). To avoid ambiguity with other scoring system,* $S^{\circ}_{ij}$ *is referred to as the* **PSM score***.*

In the rest of this article, we make the following assumption:

**Assumption 1** *The search engine provides a PSM score* $S^{\circ}_{ij}$ *of the form* $S^{\circ}_{ij} = -10\log_{10}(p^{\circ}_{ij})$ *where* $p^{\circ}_{ij}$ *is probability of a random match.*

In the setting of Ass. 1, by construction, $p^{\circ}_{ij}$ is the p-value of a test with the following null hypothesis:

$$\mathbf{H}_0^{ij} : \mathbf{spec}_j \neq \mathbf{seq}_i$$

which simply means that the peptide sequence and the observed spectrum do not correspond. A direct consequence of Ass. 1 reads:

**Corollary 1** *Under* $\mathbf{H}_0^{ij}$ (i.e. *when considering only false PSM),* $p^{\circ}_{ij}$ *is known to distribute uniformly.*

**Remark 1** *See [10] for justifications of Cor. 1.*

In other words, if symbol $\approx$ is used to represent the term "look like", then $p^{\circ}_{ij}$ corresponds to the following conditional probability:

$$p^{\circ}_{ij} = \mathbb{P}(\mathbf{spec}_j \approx \mathbf{seq}_i \mid \mathbf{spec}_j \neq \mathbf{seq}_i).$$

In practice, several spectra are acquired for each precursor ion, so that several PSMs participate to the identification of a same peptide sequence. This is why, one classically defines the **best-PSM score** of sequence $\mathbf{seq}_i$ (noted $S^{\top}_i$) as the maximum PSM score among the PSMs involving that peptide sequence:

$$S^{\top}_i = \max_{q \in [\![1,Q]\!]} S^{\circ}_{iq} \tag{2}$$

where $Q$ is the number of spectra that are possibly considered for a match onto $\mathbf{seq}_i$, and where $[\![.,.]\!]$ denotes an integer interval. Let us denote by $p^{\top}_i$ the corresponding probability, linked to $S^{\top}_i$ by Ass. 1. It rewrites as:

$$p^{\top}_i = \min_{q \in [\![1,Q]\!]} p^{\circ}_{iq} \tag{3}$$

In other words, $p^{\top}_i$ is the minimum value of a set of p-values. We would like to interpret $p^{\top}_i$ as the p-value resulting from testing of the following null hypothesis:

$$\mathbf{H}_0^{i} : \forall q \in [\![1,Q]\!],\ \mathbf{spec}_q \neq \mathbf{seq}_i$$

or with a more compact notation,

$$\mathbf{H}_0^{i} : \mathbf{seq}_i^{?}$$

the interrogation mark simply indicating that $\mathbf{seq}_i$ does not corresponds to any observed spectrum. Unfortunately, this is not possible: taking the minimum promotes small p-values, so that one should not expect the $p^{\top}$'s to distribute uniformly under the null hypothesis, which is required to have well-calibrated statistical test and to apply BH procedure. Fortunately, it is possible to recover exact calibration thanks to Prop 1.

**Proposition 1** *Let* $S_1, \ldots, S_n$ *be a set of* $n$ *scores of the form* $S_\ell = -10\log_{10}(p_\ell)$*,* $(\ell \in [\![1, n]\!])$ *where the* $p_\ell$*'s are realizations of* $n$ *i.i.d.* $\mathbb{R}_+$ *random variables,* $X_1, \ldots, X_n$*. If* $X_\ell \sim \mathcal{U}[0,1]$ $\forall \ell$*, then,*

$$Y = 1 - \left(1 - 10^{-\frac{1}{10}\cdot\max_\ell S_\ell}\right)^n$$

*uniformly distributes in* $[0, 1]$.

*Proof:*

$$\begin{aligned}
\mathbb{P}[Y \leq t] &= \mathbb{P}\left[1 - \left(1 - 10^{-\frac{1}{10}\cdot\max_\ell S_\ell}\right)^n \leq t\right] \\
&= \mathbb{P}\left[1 - \left(1 - \min_\ell \left[10^{-\frac{S_\ell}{10}}\right]\right)^n \leq t\right] \\
&= \mathbb{P}[1 - (1 - \min_\ell[p_\ell])^n \leq t] \\
&= \mathbb{P}[1 - (\max_\ell[1 - p_\ell])^n \leq t] \\
&= \mathbb{P}[(\max_\ell[1 - p_\ell])^n \geq 1 - t] \\
&= 1 - \mathbb{P}[(\max_\ell[1 - p_\ell])^n < 1 - t] \\
&= 1 - \mathbb{P}[\max_\ell[1 - p_\ell] < (1 - t)^{1/n}] \\
&= 1 - \mathbb{P}\left[\bigcup_\ell \left\{(1 - p_\ell) < (1 - t)^{1/n}\right\}\right] \\
&= 1 - \prod_{\ell=1}^{n} \mathbb{P}[1 - p_\ell < (1 - t)^{1/n}] \\
&= 1 - \prod_{\ell=1}^{n} \mathbb{P}[p_\ell \geq 1 - (1 - t)^{1/n}] \qquad (4)
\end{aligned}$$

As each $p_\ell$ is the realization of a $\mathcal{U}[0, 1]$ random variable, one has, $\forall \ell$:

$$\begin{aligned}
\mathbb{P}[p_\ell \geq 1 - (1 - t)^{1/n}] &= 1 - (1 - (1 - t)^{1/n}) \\
&= (1 - t)^{1/n} \qquad (5)
\end{aligned}$$

So that

$$\begin{aligned}
\mathbb{P}[Y \leq t] &= 1 - \prod_{\ell=1}^{n} (1 - t)^{1/n} \\
&= t \qquad (6)
\end{aligned}$$

Consequently, the cumulative distribution function of $Y$ is that of a uniform random variable. Moreover, $Y$ takes its value in $[0, 1]$, strictly. □

As well-calibration is equivalent to uniform distribution of mismatch scores, if the PSM scoring system is well-calibrated, then according to Prop. 1, the best-PSM probability can be transformed to be well-calibrated too. Concretely, uniformity under the null is thus recovered by applying a transform akin to that of Šidák correction [28] to $p^\top$, where one defines the **peptide p-value** of peptide sequence $\mathbf{seq}_i$ as:

$$p_i^\diamond = 1 - (1 - p_i^\top)^Q \qquad (7)$$

The corresponding **peptide score** (defined under Ass. 1) is noted $S_i^\diamond$.

#### S3.3 Accounting for fragmentation multiplicity

To the best of our knowledge, the peptide score resulting from Prop. 1 has never been proposed so far. However, similar mathematical derivations (*i.e.* also rooted in the Šidák correction for multiple testing) have already been applied in the proteomic context, notably to recalibrate scoring systems in a context of multiple peptide interpretations of spectra [11] – which differs from the present context. Besides, the aggregation of PSM scores into well-calibrated peptide-level scores has been already addressed by almost the same authors, notably in [14]. This article focuses on controlling the FDR at peptide-level within a TDC context, which probably explains the numerous discrepancies between their work and ours. Essentially, the article compare three different approaches: ETWO, WOTE and Fisher's combined probability test. ETWO and WOTE are both based on the best-PSM score, their difference relying in how the filtering of the non-best PSMs interplays with the TDC. As for Fisher's method, one converts all the PSM scores into PSM p-values (using Eq. 1 or a similar formula, depending on the search engine), before applying Fisher's combined probability test [29], which returns p-values at peptide level (that can finally be transformed back into peptide level scores). As a result of the comparisons, it appears that the WOTE method is the best calibrated one, tightly followed by ETWO, while Fisher method provides miscalibrated p-values. As by construction, Fisher method should provide well-calibrated p-values when combining independent tests, the authors explain this miscalibration by underlying that different PSMs pointing toward a same peptide cannot be considered as independent. We agree with this explanation and we believe it is possible to go further. Due to the the strong dependence of PSMs, using Fisher method should lead to dubious peptide scores, because of the following undesirable effect: several PSMs with intermediate scores pointing toward a given peptide are practically considered equivalent to a single PSM with an excellent score, as illustrated in the following example:

**Example 1** *Consider six spectra pointing toward a peptide sequence* $\mathbf{seq}_1$*, all with a Mascot scores of 18. Besides, one has another peptide sequence* $\mathbf{seq}_2$ *identified by a single PSM with a Mascot score of 58.084. According to Fisher method, the peptide scores of* $\mathbf{seq}_1$ *and of* $\mathbf{seq}_2$ *are equal, indicating similar confidence in both peptide identifications.*

This example contradicts with peptide scoring expectations. In fact, the PSM/peptide relationship is not the same as the peptide/protein one, so it is not surprising that Fisher method, which is helpful to switch from peptide to protein level is not to switch from PSM to peptide level.

It is interesting to note that according to our proposal, the mathematical tool suggested in [11] is of interest to solve the question raised in [14], even though bridging them has never been proposed so far. This can be explained by a noticeable drawback of the proposed peptide score: The greater the number $Q$ of PSMs pointing toward a given peptide, the smaller the score (see Figure S26). As this contradicts with the intuition that the more observed spectra, the likelier the peptide presence, a refined analysis is necessary.

From a statistical viewpoint, this penalty is well motivated: Let us consider two ions $I_1$ and $I_2$. If $I_1$ is fragmented two times more than $I_2$, it is two times more likely to reach a higher score thanks to random fluctuations (by analogy, it is easier to obtain a high score when taking the best out of 2 dices than when throwing a single dice). Thus, our Šidák-like correction is essential to avoid an increment of the peptide scores which would only result in the repetition of multiple randomized tests. Contrarily to Fisher method (discussed above through Ex. 1), the mathematical assumption underlying our correction is not that PSMs are independent; but only that their random fluctations are, *i.e.* a weaker and more realistic assumption. However, from an analytical viewpoint, this assumption is still unrealistic. In the case of high-flying ions with long elution profiles, it is possible to obtain a large number of repeated fragmentation spectra (up to 50 or 80, depending on the dynamic exclusion tuning of the mass spectrometers, on the LC length and on the complexity of the sample) with limited (and consequently correlated) random fluctuations. In such a case, the assumption on which our Šidák-like correction is based does not hold so that it should not be used to deteriorate the identification scores.

Finally, one has to find a trade-off between the necessity of correcting for multiple testing, while avoiding too systematic corrections. From our experience, such deterioration only marginally occurs and only impact ions with excellent scores which are validated regardless of a small score reduction (in fact, the higher the best-PSM score, the weaker the correction, as illustrated on Figure S27). As a result, the increased conservativeness of the correction has globally a positive effect on validation (see Results), even though more investigating for refined strategies can be of interest.

#### S3.4 Fisher combined probability test

To define the protein-level counterpart to PSMs and peptides, we leverage the intuition that fragment matches are for PSM scores what peptide matches are for protein scores. Instead of peptide sequence $\mathbf{seq}_i$, one simply has a protein sequence $\mathbf{seq}_\pi$. As for spectrum $\mathbf{spec}_j$, one has a collection of spectra $\{\mathbf{spec}_j\}_{j\in\mathbb{N}}$ that could potentially match to any of $K$ subsequences of $\mathbf{seq}_\pi$. The goal is thus to derive score and p-value for protein sequence $\mathbf{seq}_\pi$ as counterparts to peptide score and p-value (each score/p-value couple being linked by the $-10\log_{10}(.)$ transform). However, let us first make another assumption:

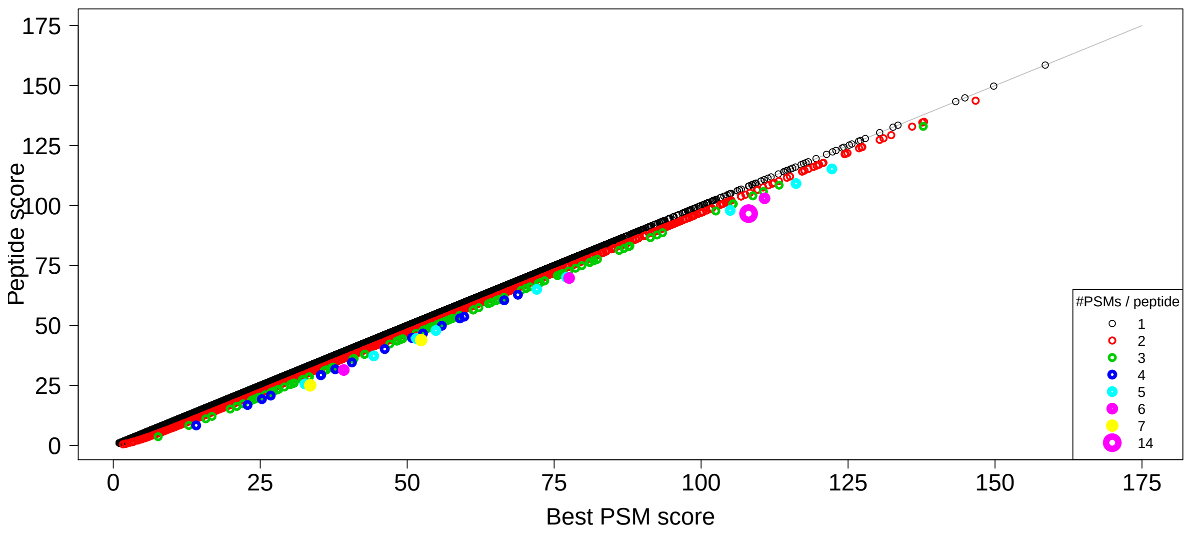

Figure S26: Peptide scores versus best-PSM scores for the *E. Coli* dataset: In the logarithmic scale induced by the conversion from p-values, the scores are deflated by a constant (proportional to the number of matches). Relatively, the confidently identified peptides are less impacted than other ones.

**Assumption 2** *The protein sequence* $\mathbf{seq}_\pi$ *does not share any subsequence with other proteins. As a result, any identified peptide sequence corresponds to a protein-specific peptide.*

**Remark 2** *Ass 2 is rather strong. In practice, one can simply restrict the analysis to peptides that are specific to a protein and discard the others, as done in many proteomic software tools. It is also classical to define equivalence classes on the peptide-protein graph, leading to so-called* ***protein-groups****.*

The next step is to define $\mathbf{H}_0^\pi$, the null hypothesis which testing would result in the desired p-value. Intuitively, if the protein $\mathbf{seq}_\pi$ is in the sample, one expects at least one spectrum to match on one of the $K$ peptide sequences; and the more matched peptide sequences, the better. However, one should not expect that all the $K$ peptide sequences are confidently matched: For example some sequences may correspond to chemical species that are difficult to ionize. Conversely, if the protein is not in the sample, only random match(es) should occur on one or few peptide sequence(s). This leads to the following null hypothesis:

$$\mathbf{H}_0^\pi : \forall k \in [\![1, K]\!], \mathbf{seq}_{\pi,k}^?$$

or

$$\mathbf{H}_0^\pi : \forall k \in [\![1, K]\!], \mathbf{H}_0^k \text{ is true},$$

where $\mathbf{seq}_{\pi,k}$ represents the $k$th subsequence of $\mathbf{seq}_\pi$. The corresponding alternative hypothesis $\mathbf{H}_1^\pi$ is that among the $K$ subsequences, at least one is matched. This reads:

$$\mathbf{H}_1^\pi : \exists k \in [\![1, K]\!] \text{ such that } \mathbf{H}_0^k \text{ is false}.$$

As a matter of fact, these null and alternative hypotheses are those of a combined probability test built on the $K$ peptide p-values $p_1^\diamond, \ldots, p_K^\diamond$. In other words, a p-value at protein level can be computed according to Fisher's method [29] (or possibly a related test, see [30, 27]), which relies on the fact that:

$$-2\sum_{k=1}^{K} \ln(p_k^\diamond) \sim \chi^2_{2K},$$

where $\chi^2_{2K}$ is the Chi-squared distribution with $2K$ degrees of freedom. This is equivalent to:

$$\frac{\ln(10)}{5}\sum_{k=1}^{K} S_k^\diamond \sim \chi^2_{2K}.$$

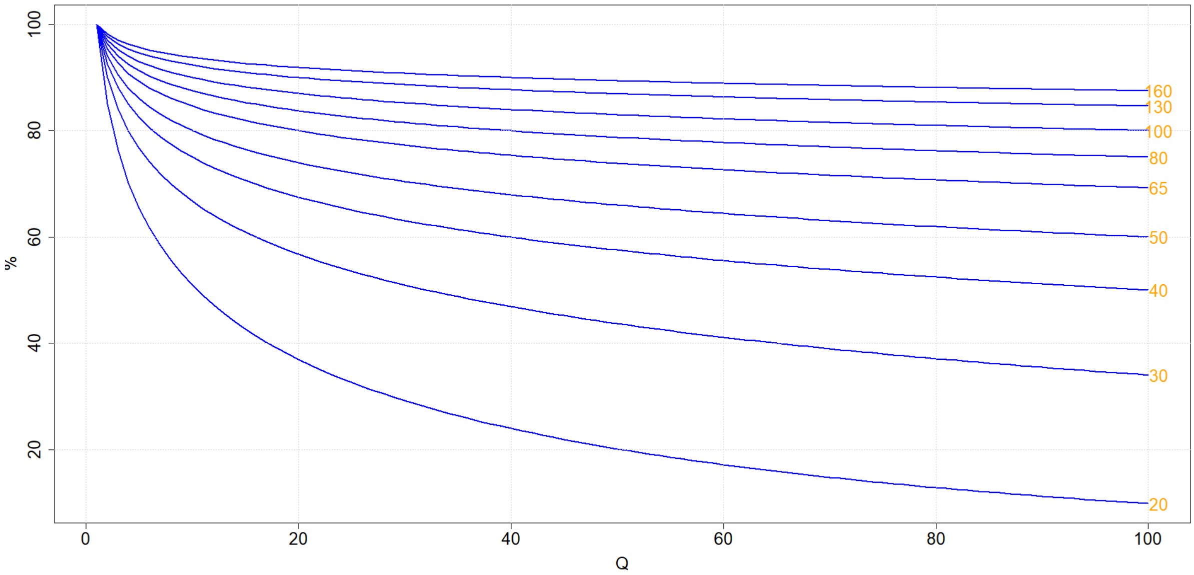

Figure S27: Peptide score expressed as a percentage of the best-PSM score in function of the number Q of PSMs: Each curve corresponds to the evolution of a given score, written in orange on the right hand side of the curve. The plot reads as follow: for Q=1, the correction is idle, so that the peptide score equates the best-PSM score (100%). The peptide score diminishes as Q grows (the plots stopped at Q=100, depicting an already extreme situation where 100 PSMs point toward the same peptide). For instance, if the best-PSM has a score of 80 (fourth curve form the top), and if reaching such a high score has required 39 additional PSMs with lower or equal scores, the peptide score is equal to 80% of the best-PSM score, *i.e.* 64.

Thus, if $f_{2K}$ denotes the density function of $\chi^2_{2K}$ and $S^{\oplus}_{\pi} = \sum_{k=1}^{K} S^{\diamond}_{k}$ (*i.e.* the sum of the $K$ peptide scores for protein $\pi$), then the p-values of the combined probability test reads:

$$p^{\dagger}_{\pi} = \int_{0.2\ln(10)S^{\oplus}_{\pi}}^{\infty} f_{2K}(x)dx \tag{8}$$

Let us call $p^{\dagger}_{\pi}$ the **Fisher p-value** and $S^{\dagger}_{\pi} = -10\log_{10}(p^{\dagger}_{\pi})$ the **Fisher score**.

#### S3.5 Enforcing the conservativeness of Fisher method

The accuracy of Fisher's method is known in the case where the $K$ p-values derive from independent tests. However, in case of dependent tests, it can possibly be anti-conservative, *i.e.* $p^{\dagger}_{\pi}$ can be underestimated (or conversely $S^{\dagger}_{\pi}$ can be overestimated), leading to too optimistic decisions. For instance, providing a too great score to a given protein makes the practitioner overly confident on the presence of the protein.

Therefore, let us analyze the possible dependencies between two tests with null hypotheses $\mathbf{H}^i_0$ and $\mathbf{H}^j_0$, respectively. In the case where protein $\mathbf{seq}_{\pi}$ is not in the sample, which corresponds to being under $\mathbf{H}^{\pi}_0$, the two tests are clearly independent: The quality of any random match is not related to the existence of other spectra matching on any other sequence of the same protein. However, if protein $\mathbf{seq}_{\pi}$ is present in the sample, the two tests should tend to reject $\mathbf{H}^i_0$ and $\mathbf{H}^j_0$, respectively: independence cannot be assumed. Therefore, the independence assumption necessary to the conservativeness of Fisher's method only holds under the null (in fact, this can be easily observed in practice: false PSMs are spread on numerous proteins while true ones tend to concentrate).

As explained above, in general, anti-conservative tests cannot be used, for they lead to too optimistic decision making. This is notably why, when using Fisher's method to combine different statistical analyses into meta-analysis, the independence of the analyses is of the utmost importance: In the case were the meta-analysis confirms the discovery of the analyses (which means rejecting their null hypothesis), it may do so with a too great confidence. This is also why, numerous alternatives to Fisher's method are available in the literature, such as for instance [31, 32].

However, in proteomics, one is seldom interested in providing a confidence level for each protein separately: Or at least, if one is, then, other tools exists, such as PEP / local FDR [24]. Most of the time, the practitioner needs to provide a list of confidently identified proteins, endowed with a quality control metric, such as the FDR. If within this list, all the Fisher scores are inflated, it is ultimately not a problem, as long as the FDR is well-controlled. In other words, it is not important to be overly optimistic with true identifications, as long as one is not with false identifications. In fact, being anti-conservative with true identifications while conservative with the false ones may be a good way to help discriminating them, and thus, reduce the FDP induced by the cut-off score (roughly, this leads the score histograms of true and false discoveries to have a smaller overlap).

As a conclusion, despite the independence assumption only holds under $\mathbf{H}_0^{\pi}$, Fisher score will not lead to an increment of false discoveries; at least, as long as one validates the protein identification list with an FDR only.

#### S3.6 Accounting for poorly identified peptides

A deeper look on Fisher's combined probability test behavior pinpoints an undesirable effect for proteomics: A very low p-value can be moderated by greater p-values. This concretely makes sense in a meta-analysis setting, where two poorly conclusive analyses will soften the conclusion of a third very conclusive analysis. However, in proteomics, two poorly reliable peptide identifications should not soften the parent protein identification, if the latter is supported by a third highly reliable (specific) peptide. Let us illustrate this on an example:

**Example 2** *Consider a protein $\pi$ with 4 peptides with score $S_1^{\diamond} = 44.09$, $S_2^{\diamond} = 1.59$, $S_3^{\diamond} = 1.59$ and $S_4^{\diamond} = 1.59$. The corresponding Fisher score is* 23.90638. *In fact, the 3 last peptides are so unreliable that they moderate the score resulting from the single observation of the best peptide. Intuitively, a score of* 44.09 *(related to highest protein evidence) would have been preferable.*

Concretely, peptides 2, 3 and 4 in Ex. 2 having small scores does not mean they are not present in the sample, but only that one did not have sufficiently good spectra.

This outlines an intrinsic limitation of Fisher score. It originates in the fact that, contrarily to the setting Fisher's method was originally designed for, mass spectrometry based proteomics relies on the Open World Assumption (OWA, [33]): an absence of observation does not mean non-existence. Concretely, a low score does not mean the peptide is not present in the sample, but only that one does not have a sufficiently good spectrum. This is why, it intuitively makes sense to consider that, given a protein, its score should not be the combination of all its peptides, but only the combination of the scores of the peptide subset which gives the highest Fisher score. This leads to the following definition of the **protein score** $S_{\pi}^{\star}$:

$$S_{\pi}^{\star} = \max_{A \in 2^{\{1,\ldots,K\}}} S_A^{\dagger} \tag{9}$$

where $S_A^{\dagger}$ denotes the Fisher score of the subset $A$ of the set of $K$ peptides that maps onto protein $\pi$. If one defines

$$p_{\pi}^{\star} = 10^{-\frac{S_{\pi}^{\star}}{10}} \tag{10}$$

then, the **protein p-value** $p_{\pi}^{\star}$ relates to the Fisher p-value by the following formula:

$$p_{\pi}^{\star} = \min_{A \in 2^{\{1,\ldots,K\}}} p_A^{\dagger} \tag{11}$$

Elaborating on methods akin to Fisher test to derive a protein level score has already been proposed in the literature. Conceptually, the method closest to ours is also the oldest [30]: The authors proposed to apply Stouffer's method [34] (which is akin to that of Fisher) on the best subset of peptides to define the protein p-value. Several differences are noticeable with respect to our proposal: First, their scoring system is used in a target-decoy context. Second, the anti-conservative behaviour induced by dependencies is not discussed. Third, the restriction to the best subset of peptides is not interpreted under the open world assumption, so that it is accompanied with a multiple test correction. Fourth, the protein score is directly based on the best-PSM score, without any intermediate recalibration at peptide level. This could lead to miscalibration, however, the specific distribution of best-PSM scores is accounted for by a Gumble law fit. More recently, numerous works have investigated a related path, yet with an objective that seems closer to the comparison and the design of protein inference rules (a subject that is not investigated in this article), rather than quality control procedures. Notably, they extensively discuss the involvement of shared peptides with regards to the protein groups, and simply resort to use TDC to estimate a protein-level FDR: In [27], authors follow a path similar to that of [30]. However, several differences exists: First, another variant of Fisher method [35] is used, which makes it possible to account for shared peptides (by down-weighting them in the combination process). Second, the method is directly applied at PSM level to derive protein-level scores. Third, it does not focus on FDR, but on individual protein-level metric instead (PEP, E-value, etc.), despite the PSM scoring system remained strongly linked to TDC. In a similar trend, works from overlapping teams ([15] and [16]) have recently investigated the use Fisher method as a protein inference rule (rather than a scoring methodology) in a TDC context, and compared it with other approaches (product of PEP, best-peptide protein surrogate, two peptide rules, etc.). Finally and concomitantly to the reviewing process of this article, a path of investigation related to that of [30] has been explored: Broadly, the authors of [36] elaborated on the best-PSM score to directly define a protein-level score (without peptide-level intermediate step) thanks to a Fisher-like method (which includes a combinatorial term accounting for multiple testing) to deliver well-calibrated p-values. However, as its predecessor, the approach is tightly attached to TDC. Investigating whether the proposed protein score could be adapted to BH framework would be an interesting future research direction.

### S4 Supporting discussion

#### S4.1 Individual vs. contextualized scores

Initially, TDC was designed to operate on raw PSM scores, *i.e.* on scores which individually quantify the similarity between an experimental spectrum and a theoretical one; irrespective of any additional information concerning the distribution of other (real or tentative) matches. However, the last decade has witnessed the publication of many "contextualized scores": Despite being rather diverse, these scores all leverage the same idea of relying on the target and the decoy score distributions to improve the discrimination between correct and incorrect matches. This can be concretely achieved by defining a delta score, *i.e.* a difference between the best PSM score and a statistics summarizing the score distribution (the second best candidate in MS-GF+and in Andromeda/Maxquant, the homology threshold in Mascot) an e-value (X!tandem) or a posterior probability (in Andromeda/Maxquant or in the Prophet post-processing tool suites). Many comparisons have experimentally assessed the improved discrimination capabilities of these thresholds. In a nutshell, by helping discarding the high scoring decoy matches, lower FDRs can be achieved.

However, it has already been noticed that these lower FDRs may not reflect the real FDP [37]. Among these contextualized scores, it is not clear which ones require a TDC, which ones require a decoy database but not necessarily a competition step, and which ones are only defined on the target match distribution. This is not necessarily an issue, as long as TDC is assumed to lead to a fair representation of the mismatch distribution. However, in this work, we hypothesized that the Equal Chance Assumption could be violated in some practical conditions. This hypothesis could indeed explain the results of [37], but it raises an issue to design our experiments: If one uses a contextualized score to assess the quality of the TDC-FDR, while the contextualized score is built on top of TDC, one faces a self-justifying loop. To cope for this, we have decided to evaluate the quality of TDC-FDR by relying on individual / raw scores, as pinpointed in [37]. The rational is the following: If TDC-FDR is stable with respect to the tuning of various mass tolerance parameters in the search engine, then, the validated PSMs with the lowest scores should have roughly the same "absolute" quality (*i.e.* irrespective of the other scores) whatever the search engine tuning. Contrarily to a well-spread belief, when one filters a PSM list at 1% FDR, it does not mean that we allow 1% of poor matches in the result. On the contrary, it means that, despite all the validated PSMs apparently depict matches of sufficient quality, 1% of them are spurious. In other words, the validated PSMs with the lowest scores are not randomly selected mismatches that make the list longer because 1% of false discoveries are tolerated; but borderline PSMs that nonetheless meet the quality standard of a 1% FDR validation. In this context, it makes sense to assume their quality should remain roughly constant whatever the search engine tuning.

Once it has been decided to use the lowest individual PSM scores to evaluate the stability of TDC across various conditions of applications, one has to select a subset of search engines to perform the experiment. This is a touchy subject as any TDC criticism can be read as a strike against a given search engine [38, 39, 40]. In our view, the five most widely used search engines are the following [41]: Andromeda (from Maxquant suite), Mascot, MS-GF+, Sequest and X!tandem. Among them, Sequest is more a core algorithm that derives in a multitude of tools [42] with different implementations, optimizations and control parameters, which ultimately lead to different identification lists. As for Andromeda, it possesses many layers of scores that cannot be accessed, and which results in behaviours that should be questioned before use in a TDC evaluation. Notably in case of very long peptides, it is customary to observe validated PSMs with fairly high posterior probability, while the similarity score is zero. Obviously, this questions the prior distributions which are involved in turning a zero score into an almost certain match, and consequently the possible construction of these posteriors from decoy matches. Finally, the last three search engines (*i.e.* X!tandem, MS-GF+ and Mascot) have already been reported to lead to similar score downfalls [40]. As a result, we have focused on Mascot (which is the most popular among the three of them and for which p-values can be straightforwardly derived) and we have postponed the study of Andromeda to Supporting Information (see Supporting Section S2.3).

As a side note, let us stress that this evaluation protocol should not be over-interpreted. Its context of use is the following, strictly: We aim at evaluating the stability of TDC, independently of the search engine or the scoring methodology. Considering, the presented evaluation protocol should not be understood as a prejudiced view on any search engine, or on any contextualized score. Notably, some contextualized score could as-a-matter-of-factly over-exploit TDC, *de facto* leading to larger FDR under-estimations (as demonstrated by [37]), while on the contrary, some others may partially cope for the problem by stabilizing the FDR. Although these questions are of interest, they stand beyond the scope of this work.

#### S4.2 Influence of the decoy database length

The TDC literature contains many references to target and decoy databases of different size (see notably Supporting Information S2.1, yet in the TDWC context). Recently, the subject was studied from a theoretical point of view, so as to provide clear guidance on how to compute the FDR in such cases. The results are as follow [43]: In principle, if $d$ (resp. $t$) stands for the number of decoys (resp. targets) that have passed the validation threshold, and $r$ is the ratio between the sizes of the whole decoy and target databases, the FDR should read

$$FDR = \frac{d+1}{r \times t}$$

However, as demonstrated in the Supporting Information to [43], this approximation only holds when $t \to \infty$ and $d/t \leq 5\%$. Consequently, when the validation threshold is set to an FDR of 1%, $r$ must be smaller than 5. Therefore, to increase $r$ in proportion to the reduction in precursor mass tolerance, the FDR should only be controlled at immaterial levels. This notably explains why, in our case, $(i)$ it was not possible to rely on an enlargement of the decoy database to recover ECA validity; $(ii)$ we had to rely on empirical null estimation instead.

[8] Andy Lin, J Jeffry Howbert, and William Noble. Combining high resolution and exact calibration to boost statistical power: A well-calibrated score function for high-resolution ms2 data. *bioRxiv*, page 290858, 2018.

[29] Ronald Aylmer Fisher. *Statistical methods for research workers*. Genesis Publishing Pvt Ltd, 1925.

[30] Victor Spirin, Alexander Shpunt, Jan Seebacher, Marc Gentzel, Andrej Shevchenko, Steven Gygi, and Shamil Sunyaev. Assigning spectrum-specific p-values to protein identifications by mass spectrometry. *Bioinformatics*, 27(8):1128–1134, 2011.

[31] Joachim Hartung. A note on combining dependent tests of significance. *Biometrical Journal: Journal of Mathematical Methods in Biosciences*, 41(7):849–855, 1999.

[32] James T Kost and Michael P McDermott. Combining dependent p-values. *Statistics & Probability Letters*, 60(2):183–190, 2002.

[33] C Maria Keet. Open world assumption. In *Encyclopedia of Systems Biology*, pages 1567–1567. Springer, 2013.

[34] Samuel A Stouffer, Edward A Suchman, Leland C DeVinney, Shirley A Star, and Robin M Williams Jr. The american soldier: Adjustment during army life. *Studies in social psychology in World War II*, 1, 1949.

[41] Kenneth Verheggen, Helge Ræder, Frode S Berven, Lennart Martens, Harald Barsnes, and Marc Vaudel. Anatomy and evolution of database search engines – a central component of mass spectrometry based proteomic workflows. *Mass spectrometry reviews*, 2017.

[42] David L Tabb. The sequest family tree. *Journal of The American Society for Mass Spectrometry*, 26(11):1814–1819, 2015.

[43] Lev I Levitsky, Mark V Ivanov, Anna A Lobas, and Mikhail V Gorshkov. Unbiased false discovery rate estimation for shotgun proteomics based on the target-decoy approach. *Journal of proteome research*, 16(2):393–397, 2016.
